# Evolutionarily conserved anatomical and physiological properties of olfactory pathway till fourth order neurons in a species of grasshopper *(Hieroglyphus banian)*

**DOI:** 10.1101/436626

**Authors:** Shilpi Singh, Joby Joseph

## Abstract

Olfactory systems of different species show variations in structure and physiology despite some conserved characteristics. We characterized the olfactory circuit of the grasshopper *Hieroglyphus banian* of family Acrididae (subfamily: Hemiacridinae) and compared it to a well-studied species of locust, *Schistocerca americana* (subfamily: Cyrtacanthacridinae), also belonging to family Acrididae. We used in vivo electrophysiological, immunohistochemical and anatomical (bulk tract tracing) methods to elucidate the olfactory pathway from the second order neurons in antennal lobe to the fourth order neurons in β-lobe of *H. banian*.

We observe highly conserved anatomical and physiological characteristics till the fourth order neurons in the olfactory circuit of *H. banian* and *S. americana*, though they are evolutionarily divergent (~57 million years ago). However, we found one major difference between the two species-there are four antennal lobe tracts in *H. banian* while only one is reported in *S. americana*. Besides, we are reporting for the first time, a new class of bilateral neurons which respond weakly to olfactory stimuli even though they innervate densely downstream of Kenyon cells.

## Introduction

Sensory systems have been well studied across insect species from many orders to elucidate principles underlying their physiology and anatomy. One of the major aims for doing so is to throw light over how the brain represents information and how it processes the information to give rise to behavior. The olfactory circuit offers a tractable and simple pathway to understand many such questions as olfaction is responsible for a range of behaviors in insects including feeding, mate selection and oviposition (Hartlieb and Anderson 1999).

One of the many insects where olfactory circuit has been well studied is the locust, *Schistocerca americana*. The olfactory information processing in *S. americana* begins when volatile molecules in the environment impinge on the ~50,000 cholinergic and excitatory olfactory receptor neurons (ORNs, Laurent 1996), housed in hair-like structures called sensilla on the antennae. The ORNs, which are the first order neurons of the olfactory pathway, transduce the chemical signals into electrical signals and transmit this information via the antennal nerve to projection neurons (PNs) and local neurons (LNs) in the antennal lobe (AL, analogous to the olfactory bulb in vertebrates, Laurent 1996; Leitch and Laurent 1996; Hildebrand and Shepherd 1997). The interaction between excitatory PNs, the second order neurons, analogous to the mitral/tufted cells of vertebrates (Hildebrand and Shepherd 1997) and inhibitory LNs, analogous to the juxtaglomerular neurons in vertebrates (Strausfeld and Hildebrand 1999), results in the olfactory stimuli being represented by an invariant spatio-temporal code spread across dynamic ensembles of active PNs (Laurent and Davidowitz 1994; Laurent 1996; Laurent et al. 1996; Leitch and Laurent 1996; Brown et al. 2005; Raman et al. 2010). The inhibition of excitatory PNs by the GABAergic LNs (Leitch and Laurent 1996; Ignell et al. 2001) intermittently, cause transient synchronization of dynamic subsets of PNs active during odor response and result in the output from the AL to be oscillatory in nature (MacLeod and Laurent 1996). The PNs, which are the only output from the AL, relay the olfactory information to the next higher centers, the mushroom body (MB, analogous to the piriform cortex in vertebrates), and the lateral horn (LH) by medial antennal lobe tract (mALT, Laurent and Naraghi 1994). The periodic synchronized output from the PNs during odor response causes oscillation in the local field potential (LFP) in the MB (Laurent and Naraghi 1994; MacLeod and Laurent 1996; Wehr and Laurent 1996), a center implicated to play a role in learning and memory (Heisenberg 2003). In addition, the dense representation of olfactory information by the AL PNs is transformed into a sparse representation in the MB by its intrinsic neurons, the Kenyon cells (Perez-Orive et al. 2002, 2004). The 50000 Kenyon cells (KCs) which are present in each MB of *S. americana* (Leitch and Laurent 1996), have a very low or near zero baseline rate and respond to odor stimuli sparsely and with high specificity (Laurent and Naraghi 1994; Perez-Orive et al. 2002; Stopfer et al. 2003; Broome et al. 2006; Jortner et al. 2007; Joseph et al. 2012). The axons of the KCs form the peduncle of the MB which bifurcates into the two output lobes-α and β (Laurent and Naraghi 1994).

The KCs output on to the β-lobe neurons (bLNs), the extrinsic neurons of MB and the fourth order neuron in the olfactory circuit (MacLeod et al. 1998; Cassenaer and Laurent 2007; Gupta and Stopfer 2014). The extrinsic neurons of the β-lobe function as a temporal channel for decoding sparse input (Gupta and Stopfer 2014).

The second termination site of the AL PNs is the LH which is thought to be analogous to the vertebrate amygdala (Friedrich 2011; Fişek and Wilson 2014). It is composed of neurons of many morphological types, a subset of which project back to the MB. The LH is implicated to play a role in innate behavior, bilateral coding, multimodal integration and concentration coding (Gupta and Stopfer 2012). Apart from these neurons, a giant GABAergic neuron (GGN) forms a critical part of this circuit. It receives its input from the MB in α-lobe and feeds it back on to the MB primary calyx. GGN acts as a normalizing agent for the MB output (Papadopoulou et al. 2011).

Many studies have compared parts of the olfactory circuit between species (Brockmann and Brückner 2001; Ignell et al. 2001; Zube et al. 2008; Rössler and Zube 2011; Bisch-Knaden et al. 2012; Jung et al. 2014; Rossi Stacconi et al. 2014; Kollmann et al. 2016; Schultzhaus et al. 2017) to emphasize the conserved principles but there is no study, to the best of our knowledge, which compares the olfactory circuit, in terms of anatomy and physiology, from the second order up to the fourth order between different species. Even though the basic plan underlying the organization of the olfactory circuit looks similar across orders of insects (Hildebrand and Shepherd 1997; Strausfeld and Hildebrand 1999; Wilson and Mainen 2006; Galizia and Rössler 2010; Martin et al. 2011), several differences (Galizia and Sachse 2010; Hansson and Stensmyr 2011) are also found right from the innervation pattern of neurons to the organization of different areas (for AL, Schachtner et al. 2005). Exhaustive data is available for comparison up to the third order of the olfactory circuit, but comparison at the fourth order level is not easy, partly because of paucity of data and partly because, we still do not know how similar a closely related species would be at this level.

In order to shed some light on this ambiguity and to thoroughly investigate the similarities and differences between two dissimilar species, we studied the grasshopper *Hieroglyphus banian* which belongs to Hemiacridinae subfamily (Dirsh 1956; Cigliano et al. 2018) and compared its olfactory circuit to a different species *S. americana* from Cyrtacanthacridinae subfamily (Kirby 1910; Cigliano et al. 2018). The two sub-families diverged from each other approximately 57 million years ago (Song et al. 2018). They are endemic to different habitats, *H. banian* is native to the Indian subcontinent and Vietnam and *S. americana* to North America (Cigliano et al. 2018) and they differ in their major host plant preferences as well. *H. banian* is a major pest of paddy apart from consuming a wide variety of food plants from Poaceae and Cyperaceae families (Das et al. 2002; Mandal et al. 2007). On the other hand, *S. americana* feeds on citrus plants, corn, soybean, bean and several species of grasses (Capinera 1993; Squitier and Capinera 1996). Additionally, *H. banian* has a single generation in a year (Mandal et al. 2007) while *S. americana* has two generations per year (Kuitert and Connin 1952). As the olfactory circuit plays a major role in all these behaviors, including feeding and oviposition, we, therefore expect to find differences between the olfactory circuits of the two species.

To compare the olfactory circuit anatomically and physiologically between the locust *S. americana* and *H. banian*, we recorded responses of neurons to odor stimuli intracellularly from different levels (AL PNs and LNs, MB KCs, LHNs, GGN and bLNs) and filled them with dyes post-recording to characterize their morphologies. We also recorded extracellularly from the MB simultaneously to study the characteristics of local field potential during odor response. Additionally, we injected tract tracing dyes in different areas to determine anatomical connectivity between them.

Our results show that the anatomy and physiology of the neurons in the olfactory pathway in the two species are highly conserved till the fourth order. However, we found a major difference in the number of antennal lobe tracts (ALTs) to MB in *H. banian,* when compared to that reported in *S. americana*. There is only one reported ALT, the medial ALT in *S. americana* (Laurent and Naraghi 1994), while our tract-tracing results show that in *H. banian*, there are 4 ALTs-medial ALT, transverse ALT, lateral ALT and mediolateral ALT. We have also discovered a new class of bilateral mushroom body extrinsic neurons (MBEN) in the circuit. These neurons have dense innervations in the olfactory area, downstream of Kenyon cells of MB, but are surprisingly, weakly responsive to odor stimuli unlike other fourth order neurons, the bLNs and GGN.

## Materials and methods

### Animals

Adult grasshoppers (*Hieroglyphus banian*) of either sex from a crowded colony bred in-house at the Center for Neural and Cognitive Sciences, University of Hyderabad were used for the experiments. Animals were reared at relative humidity: 70%; temperature: 29°C and 14L: 10D photoperiod.

### Animal preparation for experiments

All the experiments were performed on unanaesthetized animals at room temperature (25°C). The animals were restrained in a plastic platform using tape after the limbs and wings were removed. This was done so that there was no movement to disrupt the recording process. The plastic platform was fixed onto a clay stage placed on a petri-plate. Both the antennae were threaded through a thin PTFE tubing to restrain it during recording. A wax cup was built around the head of the animal to hold the insect physiological saline (140 mM NaCl, 5 mM KCl, 5 mM CaCl_2_, 4 mM NaHCO_3_, 1 mM MgCl_2_, 6.3 mM HEPES, pH 7.1; Laurent and Naraghi 1994) which superfused the brain throughout the experiment. To expose the brain, the cuticle, fat bodies and air sacs above it were removed. The gut was removed from an incision made at the posterior end which was ligatured afterwards. A small platform made from wire and thinly covered with wax was positioned carefully below the brain to elevate it and to give additional stability during electrophysiological recordings. The brain was then desheathed using protease (P-5147, Sigma) to remove the protein sheath covering it and to access the neurons.

### Odor delivery

Odor stimuli were delivered to the antennae by the means of PTFE tubing, as described previously (Brown et al. 2005). Odors were delivered by injecting a measured volume (~0.1 L/min) of the static headspace above the odorants into a stream of desiccated air (~0.75 L/min). The desiccated air was projected continuously on to the antennae by a tube kept 3 cm away from it. The delivery of the odorants was controlled by a valve and after the delivery a large vacuum pipe (16 cm diameter) positioned behind the antennae removed the odorants, to prevent it from lingering in the airspace. The odorants (5 ml of each) were kept in 30 ml glass bottles and used at 100%, 10% and 1% dilution (diluted in mineral oil v/v). The odorants used in this study were 1-Hexanol (hex), 2-Octanol (oct2ol), Octanoic acid (octac), Geraniol (ger), 1-Octanol (oct1ol), mineral oil (MO) and fresh wheat grass. All the odorants and mineral oil were procured from Sigma-Aldrich, India.

### Field emission scanning electron microscopy (FE-SEM)

FE-SEM (Zeiss Ultra 55) was done for the different types of sensilla present on the antennae of *H. banian.* As outlined by Ochieng et al. (1998), the antennae were cut near the base and immediately immersed in 70% ethanol at room temperature for fixation. After three days, the antennae were dehydrated using an ascending ethanol series of 80%, 90% and 100% for 30 minutes each. The antennae were stored in 100% ethanol at 4°C till the day of imaging. Half an hour before imaging, the ethanol was pipetted out from the eppendorf containing the antenna and it was dried using hot air dryer for 2 minutes. The antennae were subsequently air dried for 15 minutes. The antennae were sputter coated with gold palladium for 90 seconds and the imaging was done at 3/20 kV EHT in FE-SEM (Zeiss Ultra 55).

### Antennal backfills

The antenna of an animal mounted on a petridish, as described above, was cut till the pedicel and dextran biotin (D-7135, Invitrogen) was inserted into the stump. It was then covered with Vaseline to avoid desiccation. The mounted animal was kept at 4°C overnight and the brain was dissected out the next day and processed as those for neural tract tracing described below.

### Neural tract tracing

To ascertain the anatomical connectivity between different areas and the calyx of MB, we injected dextran biotin (D-7135, Invitrogen) or dextran tetramethyl rhodamine (D3308, Invitrogen) either in the MB calyx, AL or β-lobe in different preparations. The brains were continuously perfused with insect physiological saline and dissected out after six hours. They were fixed in 4% paraformaldehyde for four hours, washed in phosphate buffered saline (PBS; 3X30 minutes) and subsequently transferred to 3% triton-X in PBS (PBST) for one hour, for permeabilizing the membrane. The tissues were incubated at 1:1000 dilution with streptavidin-conjugated Alexa Fluor 488 or 568 or 633 (S11223 or or S11226 or S21375, Invitrogen) for five days with intermittent shaking. After five days, the brains were washed in PBST (3X20 minutes) and then with PBS (3X20 minutes). Thereafter, the brains were dehydrated by running it through an ascending ethanol series (30%, 50%, 70%, 80%, 90% and 100%; 20 minutes in each) and finally they were mounted in methyl salicylate (M-6752, Sigma Aldrich, India). The whole mounts of brains thus obtained were imaged using confocal laser scanning microscope (CLSM).

The brains with cells filled with intracellular dye injection were dissected out, fixed in 4% paraformaldehyde for four hours and processed as described above before imaging using CLSM.

### Immunohistochemical study

Immunocytochemistry was done on brains with dye-filled cells, for GABA and the synaptic density protein bruchpilot, using rabbit polyclonal anti-GABA (Sigma, A2052) and mouse monoclonal nc82 (DSHB, Iowa city, IA; donated by E. Buchner) primary antibodies respectively. The brains were fixed in 4% paraformaldehyde for four hours, washed in PBS (3X30 minutes), permeabilized using PBST for one hour, and then transferred to 10% normal goat serum in PBST for blocking. They were then incubated with the primary antibodies, rabbit anti-GABA (1:1000) and mouse nc82 (1:1000) for five days at 4°C. After five days, the brains were rinsed in PBS (3X20 minutes) and transferred to PBST containing either goat anti-rabbit/anti-mouse Alexa Fluor 488 or 568 or 633 IgG secondary antibody (Invitrogen, A-11008 or A-11011 or A-21070) at 1:1000 dilutions. After five days of incubation in secondary, they were first washed in PBST and then PBS (3X20 minutes each), ran through an ascending ethanol series for dehydration and mounted in methyl salicylate. The tissues were imaged using CLSM.

### Confocal microscopy and image processing

The whole mounts of brains were imaged using confocal laser scanning microscope (Leica TCS SP2, Leica Microsystems or Zeiss LSCM NLO 710) with an objective of 10X or 20X. The tissues were scanned at a resolution of 512X512 or 1024X1024 pixels respectively. The intensity and gain of the lasers were manually adjusted. The confocal stacks were later processed using public domain software Fiji ImageJ 1.47v, Inkscape 0.48 and GIMP 2.8. The images were altered only for brightness and contrast.

In few cases (Figs. 6 and 9), the dye-filled neurons, when not clear while using z-projection, were traced using the simple neurite tracer plug-in (Longair et al. 2011) in Fiji ImageJ (National Institute of Health, Bethesda, MD).

### Electrophysiology

#### Extracellular recordings

LFP was measured from the cell body layer and calyx of MB using custom made blunt borosilicate glass microelectrodes (impedance <10MΩ after filling with saline), pulled using a horizontal puller (P97; Sutter instrument, Novata, CA). The signal was amplified (1000X) using Axoclamp 900A (Molecular Devices) amplifier and it was low pass Bessel filtered at 80 Hz. Subsequently, the amplified signal was digitized and acquired at a sampling rate of 10 kHz using Clampex 10.3 software and a Digidata1440A interface (Molecular Devices).

#### Intracellular recordings

Intracellular recordings were done using custom made sharp borosilicate glass microelectrodes (impedance 60-200 MΩ after filling 0.2 M KCl or an intracellular dye solution made in 0.2 M KCl) from antennal lobe projection neurons and local neurons, Kenyon cells, β-lobe neurons, lateral horn neurons and giant GABAergic neuron. The signals were amplified (10X/5X) using Axoclamp 900A (Molecular Devices) amplifier and low pass filtered at 4 kHz. Subsequently, the signal was digitized at a sampling rate of 10 kHz using Clampex 10.3 software and a Digidata1440A interface (Molecular Devices). Intracellular staining of neurons after recording from them was done by filling the neurons ionotophoretically with 2% Neurobiotin (SP-1120, Vector Labs) by injecting 1-4 nA of current at 2-5 Hz pulse for 30-60 minutes.

### Data analysis

All electrophysiological data were analyzed offline using custom programs in MATLAB (MathWorks).

## Results

### Types of sensilla on the antenna and major neuropils of the brain in *H. banian*

The antenna of *H. banian* (Fig. 1a) is characterized by different types of sensilla present on its surface (Fig. 1b). A sensillum, putatively mechanosensory, longer and narrower as compared to the olfactory sensilla, was observed at the basal segment of the antenna (Fig. 1c). It did not have pores on its surface. Field emission scanning electron microscopy (FE-SEM) of the antennae showed the presence of four types of olfactory sensilla: basiconica, chaetica, coeloconica and trichodeum, (Fig. 1d-g). The basiconic sensillum (Fig. 1d), which is set in a shallow depression in the cuticle of the antennae, is characterized by numerous pores on its surface while sensillum chaeticum (Fig. 1e) is characterized by longitudinal grooves on the cuticular surface. Sensillum coeloconicum (Fig. 1f) is found in pits present on the antennal surface and has longitudinal ridges on its surface. It is shorter than other sensillar types. Sensillum trichodeum (Fig. 1g) has fewer pores on its surface when compared to basiconic sensillum. The density of olfactory sensilla is high at the tip and lowest at base, in agreement with the variation of the strength of electroantennogram (EAG, Supplementary Fig. A1) recorded from the tip and base of antennae.

**Fig. 1.**
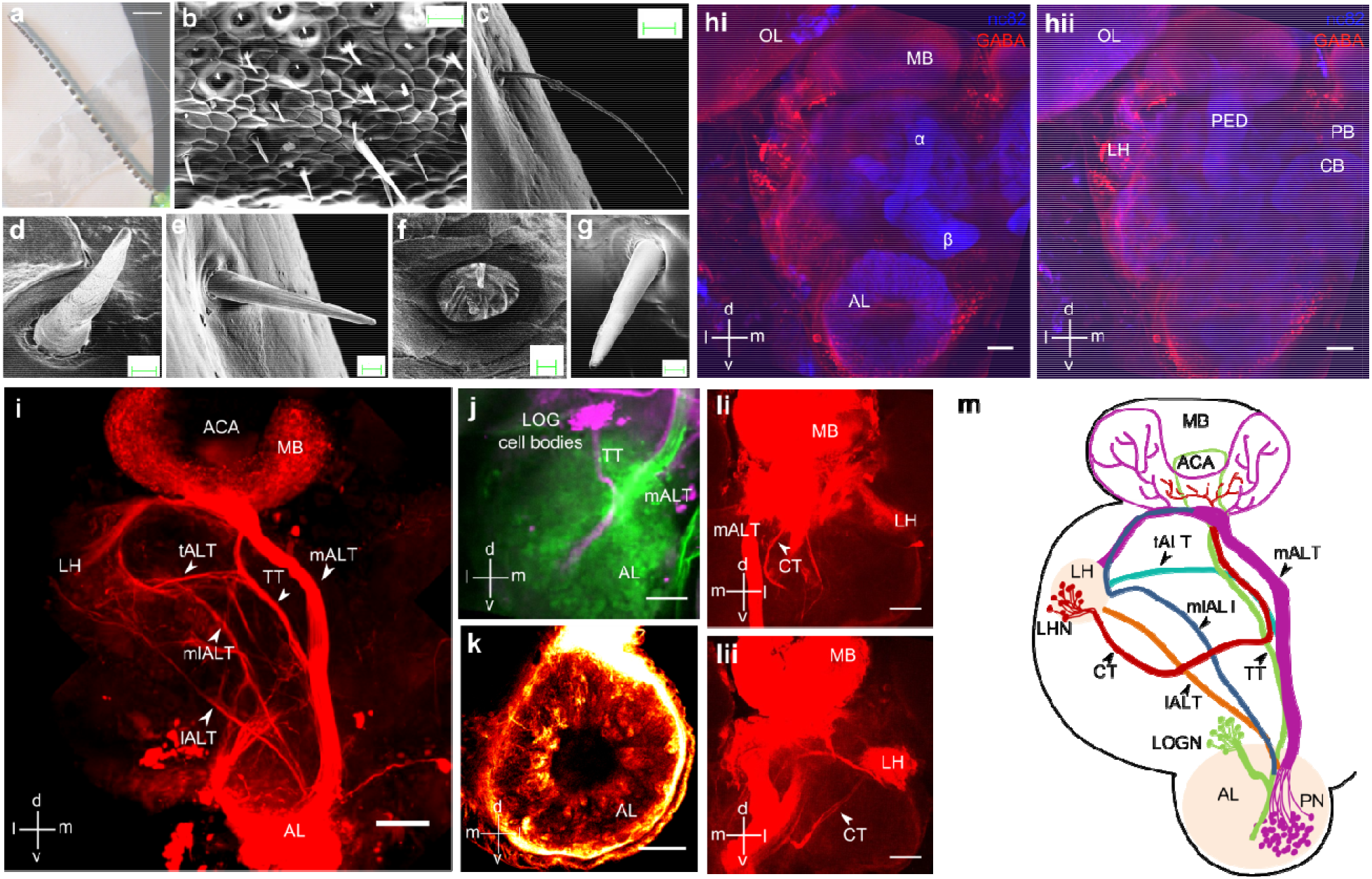
First and second order structures of the olfactory pathway in *H. banian*. **(a)** Antenna of *H. banian*. **(b-g)** FE-SEM micrographs of different types of sensilla present on the antenna **(b)** Sensilla of different types present on a segment of antenna. **(c)** A mechanosensory sensillum, present at the basal segment of the antenna **(d)** Sensillum basiconicum, with numerous pores on its surface **(e)** Sensillum chaeticum, with longitudinal ridges on its surface **(f)** Sensillum coeloconicum, located in a pit on the antennal surface and characterized by longitudinal grooves on its surface. It is the shortest among all sensillar types. **(g)** Sensillum trichodeum. This also has pores on its surface but the density is less as compared to S. basiconicum. **(hi)** and (**hii)** Major neuropils visualized by immunohistochemistry against pre-synaptic density protein bruchpilot (nc82, blue) and anti-GABA (red) antibody. The clearly discernible neuropils are mushroom body (MB) and its pedunculus (PED), antennal lobe (AL), lateral horn (LH), optic lobe (OL), protocerebral bridge (PB) and central body (CB), the last two form part of the central complex. The MB bifurcates into two lobes: α-lobe and β-lobe **(i)** Dye fill from the AL shows multipl antennal lobe tracts (ALTs) originating from the AL and projecting to the higher olfactory regions-MB and LH. The major olfactory tract, medial ALT (mALT), projects to MB calyx and LH while the lateral ALT (lALT) and the transverse ALT (tALT) project only to the LH. The mediolateral ALT (mlALT) projects to the LH first and then the MB calyx. The tritocerebral tract (TT), which originates from the cell bodies of lobus glomerulus neurons (LOGN) in tritocerebrum, runs along the mALT and separates from it midway, projecting to the accessory calyx (ACA) in MB. **(j)** The cell bodies of the LOG which are located dorsolaterally near the AL, give rise to the TT which runs ventrally for some distance before bifurcating-one branch runs ventrally, while the other leaves the AL at the same point as the mALT running along its lateral edge. **(k)** Dextran biotin fill from the antennae shows the microglomerular nature of innervations of ORN terminals in AL **(l i)** and **(l ii)** Dye fill from the calyx of MB shows a curved tract (CT, arrowhead), not reported till now in any insect species. This tract arises from the LH cell bodies and terminates in the MB. (**m**) Schematic of the AL tracts, made after combining the images in **i, j, li** and **lii.** Images **(b-g)** are from the same animal. Images **(h-l)** are from different animals. Scale bars: 2mm **(a),** 20 μm **(b, c),** 2 μm **(d, f, g),** 3 μm **(e),** 100 μm **(h-l)**

The whole brain of *H. banian* was immunostained for the synaptic density protein bruchpilot and GABA using nc82 and anti-GABA primary antibodies respectively. nc82 stained the major neuropils found in insect brain, namely, antennal lobe, optic lobe, mushroom body calyx, peduncle of mushroom body, α-lobe, β-lobe and central complex (protocerebral body and central body), while anti-GABA antibody stained the GABA-positive cell bodies (Fig. 1hi and hii).

### New AL tracts discovered in *H. banian*

Dye fills from the AL showed us three new tracts previously unreported in any species of family Acrididae (order: Orthoptera), in addition to the major tract, mALT. The three new tracts observed are transverse antennal lobe tract (tALT), mediolateral ALT (mlALT) and lateral ALT (lALT) (Fig. 1i). We have used the same terminology for these new tracts as specified by Ito et al. (2014)because of their similarities to the AL tracts known in other species. The mALT exits the AL dorsomedially, projects to the MB calyx first and then terminates in the LH, its trajectory consistent with that found previously in Orthopterans (Ignell et al. 2001). tALT and lALT project to the LH directly, bypassing the MB completely. tALT runs along the mALT and deviates from it midway, travelling laterally to end in LH. lALT exits the AL adjacent to the exit point of mALT, but turns dorso-laterally to become the lateral-most AL tract and terminates in the LH. The mlALT, on the other hand, projects both to LH and MB calyx, but in the opposite order as compared to mALT. Though it exits the AL at a location lateral to the mALT, it crosses the lALT and runs in the medio-lateral position to the LH first. It then turns dorsally to run further and terminate in the MB calyx.

We also observed the tritocerebral tract (TT), from the lobus glomerulus (LOG) to the accessory calyx of the MB (Fig. 1i and j). The LOG receives its input from the ORNs present on the maxillary palps of the insect (Ernst et al. 1977). The TT is formed by the axons of the LOG cell bodies which are located dorso-laterally of AL, at the boundary of deuto- and protocerebrum and adjacent to the entry point of the antennal nerve (Fig. 1j). It travels ventrally for a short distance and then bifurcates; one branch runs laterally along the mALT and exits the AL from the same place as mALT. After travelling along the mALT for some distance, it separates from it midway and terminates in the accessory calyx of MB.

Dye fill from the antenna shows the ORN terminals and the microglomerular nature of AL (Fig. 1k). A central area is seen which is devoid of ORN innervation (dark in the figure). This area is surrounded by bright radial fibers of ORNs and bright spots indicative of microglomeruli.

We also observed a novel tract, curved in shape and named as such (Curved Tract, CT), which connects the LH and MB. (Fig. 1 li and lii). This tract was visible in five separate preparations in which dye was injected in the MB calyx. We could see the complete tract only in one preparation but even in partially filled preparations, it could be recognized unambiguously. From our data, we saw that the CT starts from the cell bodies in LH and runs ventro-medially till it reaches near the peduncle of MB. It then turns medially and runs towards the mALT, traversing just below the point at which the peduncle bifurcates into the two lobes. It turns again to run dorsally along the mALT, and terminates in the calyx at a location between the peduncle and the mALT. The CT has not been reported till now, to the best of our knowledge and further study is required to elucidate its role in the circuit.

### Second order neurons of AL

We found two morphological types of neurons in the antennal lobe of *H. banian,* which is consistent with previous findings in *S. americana* and Orthoptera in general (Laurent 1996; Ignell et al. 2001; Galizia and Rössler 2010). One type corresponds to the multiglomerular projection neuron (PN) found in the AL of *S. americana*, the axon of which leaves the AL and projects to the calyx of MB and LH. The second type is the inhibitory local neuron (LN), whose arborization is restricted to the AL.

Intracellular recordings from a PN showed that it responds to all odorants tested with different spatio-temporal patterns of activity (Fig. 2a). We were able to fill two PNs in *H. banian,* both of which were multiglomerular. One of the fills is shown in Fig. 2b. Anti-GABA immunohistochemistry of this PN showed it to be GABA-negative (Fig. 2b). Additionally, different PNs responded to the same odorant with different temporal patterns of activity (Fig. 2c). The odor responses of all the PNs were temporally patterned with odor-cell specific sequence of excitations, inhibitions and quiescence. All PNs recorded showed spontaneous spiking activity at baseline.

**Fig. 2.**
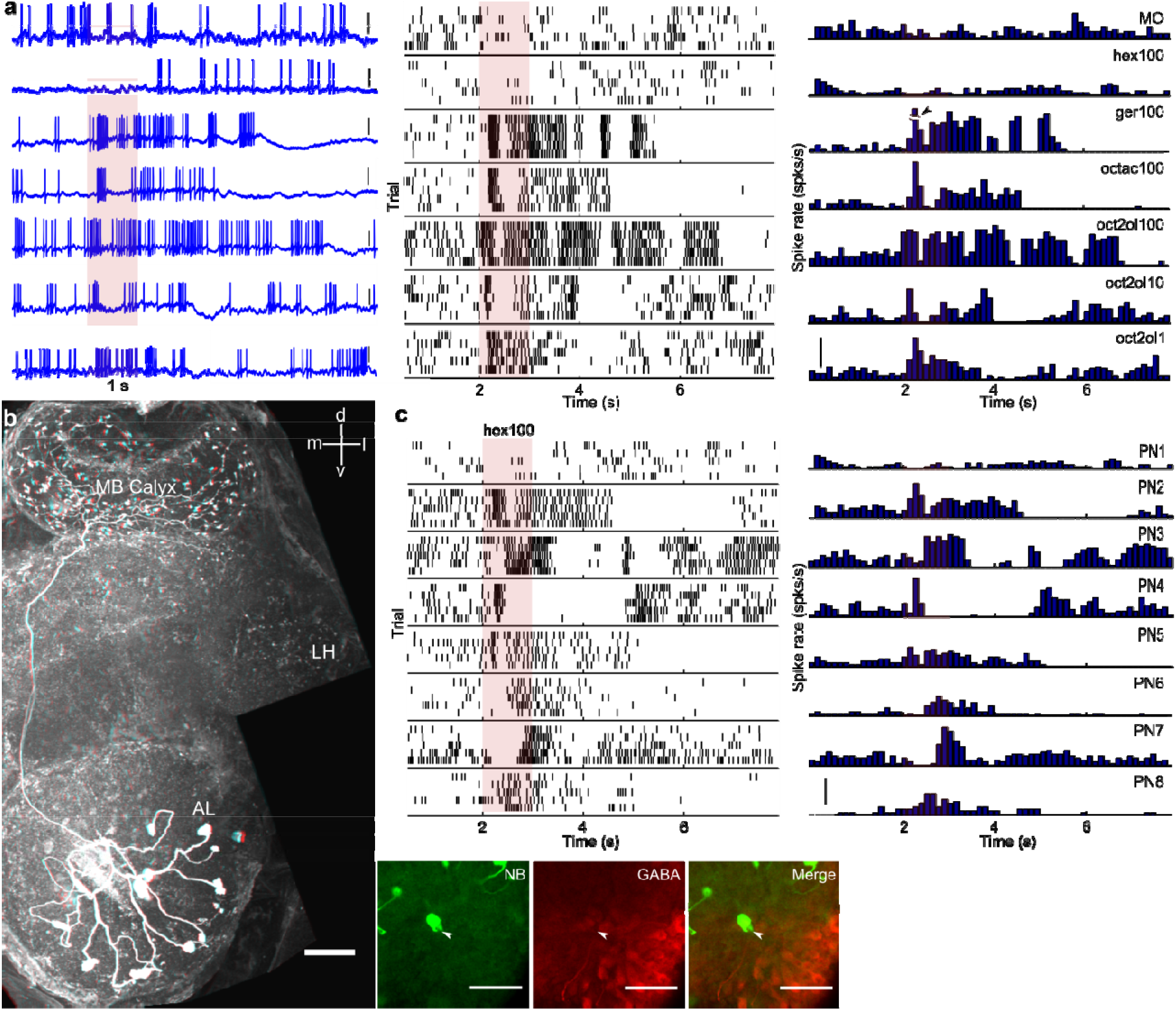
Morphology and odor response of antennal lobe (AL) projection neurons (PNs) in *H. banian*. **(a)** Odor response of an AL PN to different odorants. The same PN responds to different odorants with different temporal patterns consisting of epochs of excitation, inhibition or quiescence. MO: mineral oil, hex100: 1-Hexanol 100%, ger100: Geraniol 100%, octac100: octanoic acid 100%, oct2ol100/10/1: 2-Octanol100 %/10%/1%. Single traces are shown first, followed by rasters in the middle panel and peri-stimulus time histogram (PSTH). Scale bar: 5 mV (raw traces); 20 spks/s (PSTH); arrowhead: ~62 spks/s **(b)** Intracellular fill of AL PN shows that this PN in *H. banian* has multiglomerular arborization in AL. The axon of this PN leaves the AL dorsally and arborizes in the MB calyx covering a large area. It continues further before terminating in the LH. Anti-GABA immunohistochemistry (inset) of this PN shows that it is non-GABAergic. Scale bar: 100 μm **(c)** Odor responses of eight different PNs to the same odorant shows that different PNs respond with different patterns of activity to the same odorant. Scale bar: 20 spks/s (PSTH)

We also filled a LN in *H. banian*, and tested it for GABA immunoreactivity (Fig. 3a). This LN arborizes throughout the AL and tested positive for GABA. GABA immunostaining of the whole brain also revealed a cluster of GABA-positive cell bodies in the AL (Fig. 3b). Whether this cluster of GABA-positive cell bodies belongs only to LNs (as reported in *S. americana,* where all LNs are inhibitory and all PNs are excitatory (Leitch and Laurent 1996) or whether GABA-positive PNs are also present (as reported in *Drosophila and Apis mellifera*; Schäfer and Bicker 1986; Okada et al. 2009) needs to be investigated further. In addition, whether all LNs in *H. banian* are GABA-positive, needs to be thoroughly explored too. Intracellular recordings from different putative LNs show that unlike the PNs, they respond to odor stimuli with non-uniform small-amplitude spikes, putatively calcium-mediated. They too have a baseline spontaneous activity similar to the PNs, and show multiphasic odor responses (Fig. 3c).

**Fig. 3.**
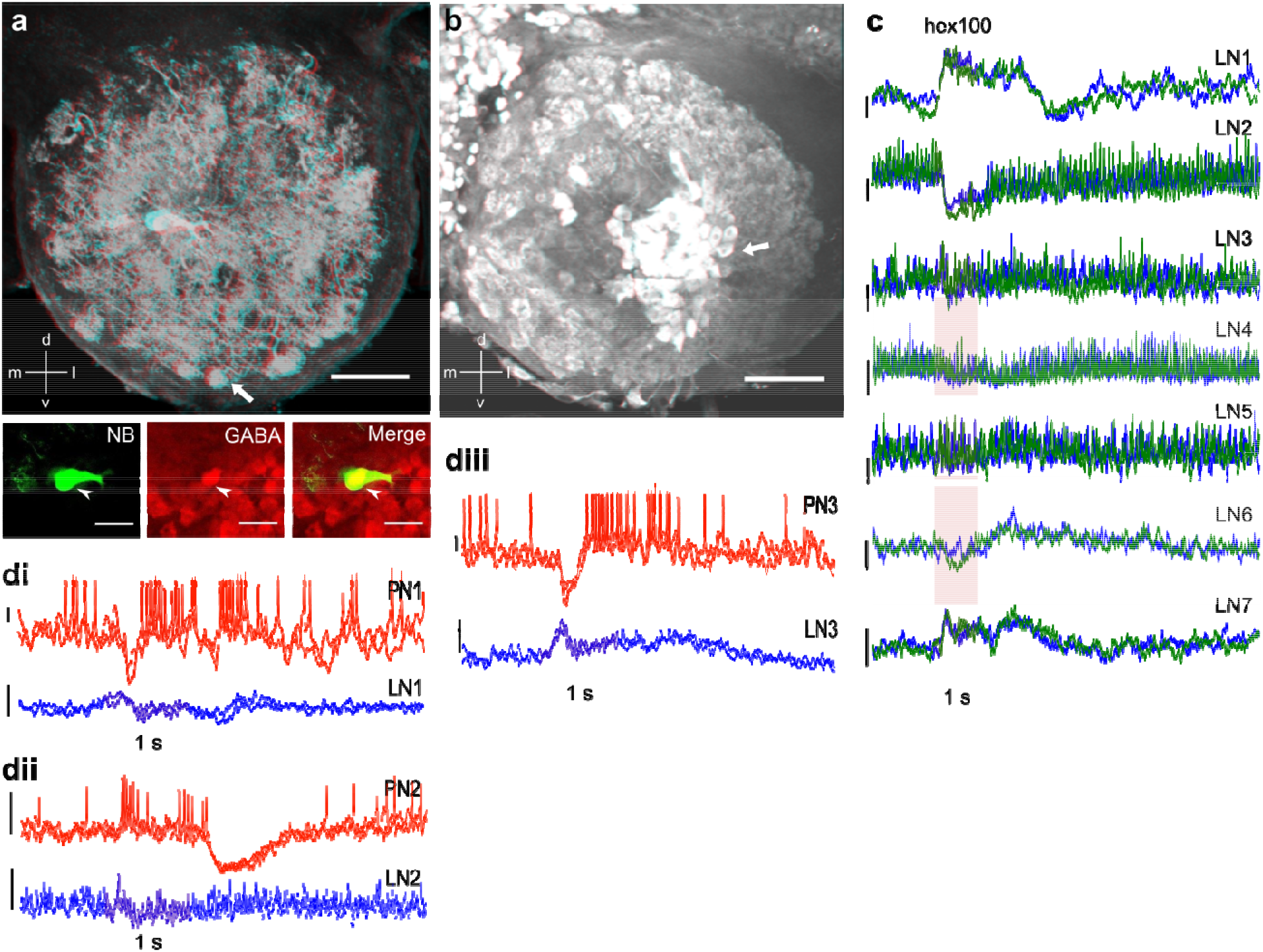
Morphology and odor responses of AL local neurons (LNs) in *H. banian*. **(a)** Intracellular fill of an AL LN showing its dense arborizations throughout the AL. The dense clusters (arrow) are indicative of microglomeruli of the AL. Scale bar: 100 μm. Anti-GABA immunohistochemistry of the AL shows that this LN is GABA-positive (inset). Scale bar: 50 μm. **(b)** Immunohistochemistry of the whole brain also reveals a GABAergic cell cluster (arrow) in the AL, putatively LNs. Scale bar: 100 μm **(c)** Odor responses of seven different non-spiking cells, putatively LNs, show the different kinds of multiphasic responses to the same odorant. Two trials (blue and green) of each LN are superimposed on each other to show the consistency of their odor responses. LN1 shows spikelets riding on top of depolarization of the membran potential, in response to odor stimulus while LN2 shows hyperpolarization for the same. LNs 3, 4 and 5 show weak odor response. LN6 shows an off response. LN7 shows depolarizing response followed by hyperpolarization of the membrane potential. Scale bar: 20 mV **(di-diii)** Paired recording of antennal lobe interneurons (PN and LN). Reliable occurrence of correlated or anti-correlated changes in the odor responses of PN-LN pairs indicates the possibility of interactions between them. Scale bar: 2 mV

Paired recordings between three different PN-LN pairs showed co-variation of activity in these cell pairs, implying possible synaptic connections between them (Fig. 3di-diii). The LN in the first and third pair (Fig. 3di and 3diii) depolarizes during odor response which is reflected in the inhibition of the PN after a delay, indicating a possibility of LN to PN synaptic interaction between them. The second PN-LN pair (Fig. 3dii) doesn’t seem to have correlated change in odor response, possibly not having a synaptic connection.

### Third order higher olfactory centers-MB and LH

We recorded the LFP from the primary calyx and the cell body layer of the MB and are reporting here for the first time that they are negatively correlated to each other during the baseline as well as during odor response (Fig. 4a). The LFP recorded shows characteristic ~25 Hz oscillations during odor response and the power of the oscillations (between 15-40 Hz) increases over repeated trials as reported in *S. americana* and *Manduca sexta* (Stopfer and Laurent 1999; Ito et al. 2009). We also show that the power of the lower frequencies (1-5 Hz) decreases over repeated trials, concomitant with an increase in power at higher frequencies (between 15-40 Hz; Fig. 4b).

**Fig. 4.**
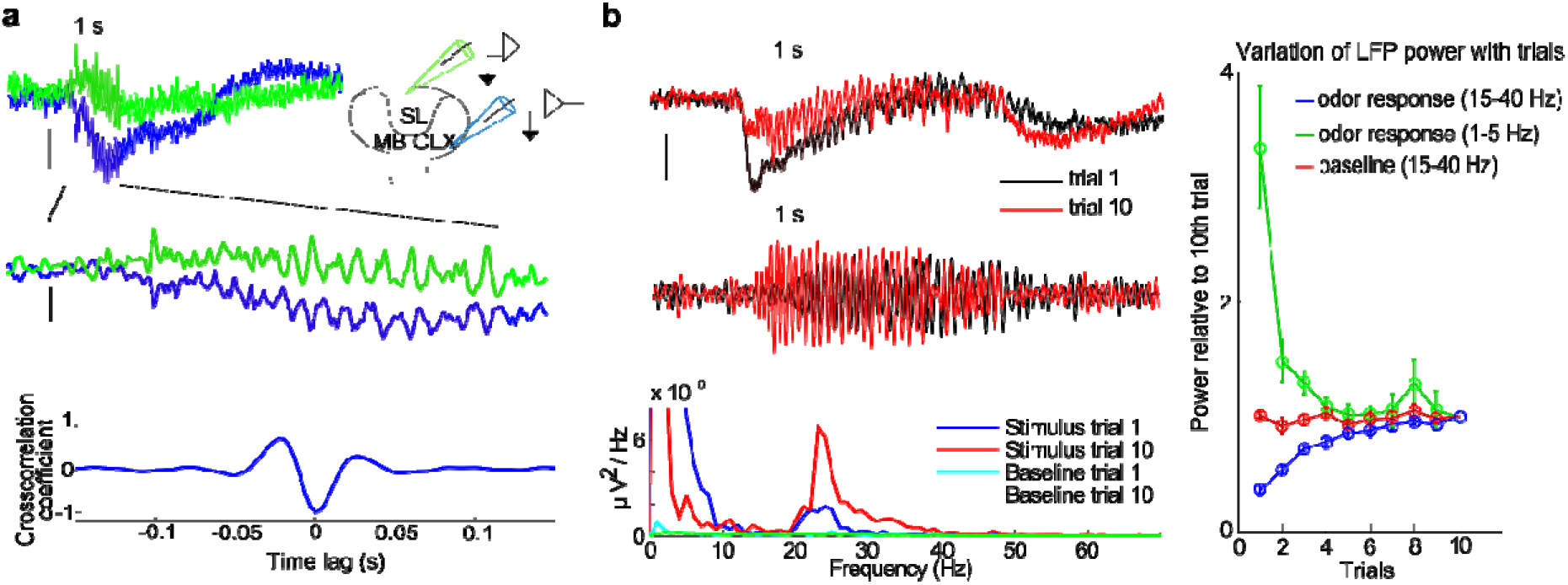
Characteristics of the local field potential (LFP) recorded from mushroom body (MB) during odor response. **(a)** LFP recorded simultaneously from the soma layer (SL, green trace) and calyx of MB (MB CLX, blue trace) show oscillations during odor response which are negatively correlated and out of phase to each other. Schematic shows the recording locations of the two electrodes in the MB. Scale bar: 100 μV **(b)** The strength of the LFP oscillations during odor response recorded from the calyx of MB increases over repeated trials. The top panel is the raw data (Scale bar: 0.5 mV). It is filtered at 15-40 Hz and shown in the middle panel (Scale bar: 0.2 mV). The third panel shows the power spectrum of the LFP during odor response, with a clear peak at ~25 Hz, the predominant frequency in the signal. The right panel shows the chang in the strengths of the component frequencies during odor response, over the course of ten trials. The lower frequencies (1-5 Hz, green trace) show a decrease in their strength over repeated trial whereas the frequencies between 15-40 Hz (blue trace) increase in strength from the first to the tenth trial

We recorded from the calyx of the MB using electrodes with high impedance to target the Kenyon cells’ dendrites. Kenyon cells show subthreshold membrane oscillations during odor response (Fig. 5a). Cross-correlogram computed between the LFP recorded from MB calyx and KC membrane potential show that they are periodically synchronized (Fig. 5b). Recordings from four different KCs show that their baseline spiking rate is close to zero (Fig. 5c).

**Fig. 5.**
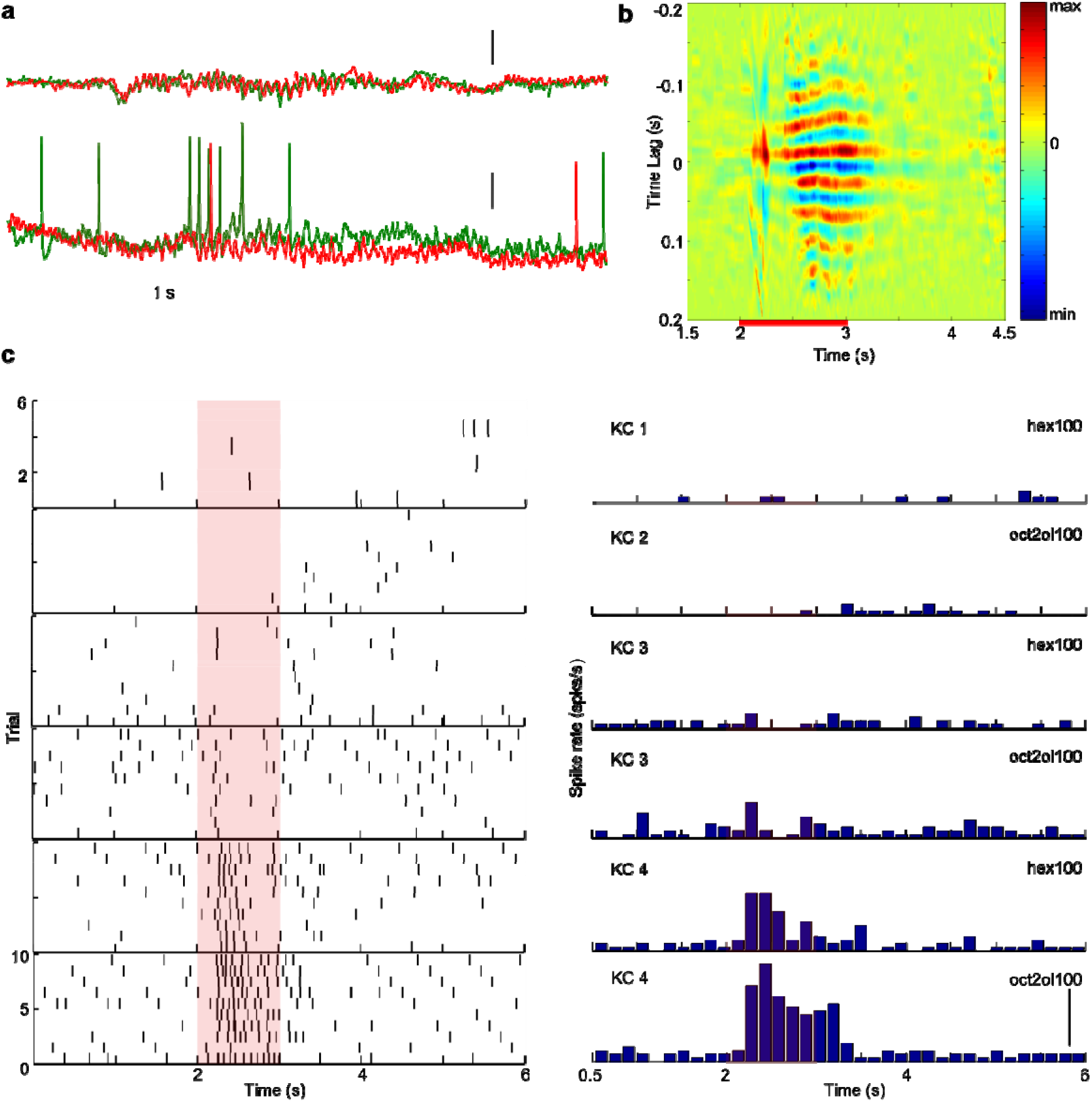
Odor response of Kenyon cells (KCs), the intrinsic neurons of MB. **(a)** Intracellular recording from KC in MB shows subthreshold oscillations during odor response. The upper trace shows the LFP recorded from the MB calyx with oscillations during odor response. Scale bar: 200 μV (LFP), 1mV (KC) **(b)** Cross-correlogram computed between LFP recorded from MB calyx and intracellular recording from KC shows that the KC membrane potential during odor response is synchronized with the LFP **(c)** 4 KCs recorded from 3 different animals show that they have a low baseline rate and they fire in a specific way in response to odor. Scale bar: 10 spikes/s

The LH is a synaptic area located in the lateral protocerebral lobe and receives major input from the antennal lobe projection neurons (Ernst et al. 1977). This synaptic area along with the LH neurons’ (LHNs) cell bodies (arrowhead) can be seen in dye fills from the MB calyx in *H. banian* (Fig. 6a). Recently, the LH area, which was previously thought to be an unstructured neuropil, was shown in moths to have a distinct structure of two linked toroids at a particular depth (Ian et al. 2016). But we could not discern any such organization of the LH in *H. banian*. We also recorded intracellularly from neurons in this area and filled six of them, each with a distinct morphology (Fig. 6 bi-gi). Few of them had dense arborization in the LH (Fig. 6 ci, fi, gi) while others had sparse arborization (Fig. 6 bi, di, ei), but all of them showed clear odor response (Fig. 6 bii-gii). The pattern of activity of all the LHNs consisted of an ongoing baseline spike rate that increased during odor response with subthreshold membrane oscillations. Only one LHN (Fig. 6 gi), out of the six, which we successfully filled in *H. banian,* looks morphologically similar to the one reported in *S. americana*, where 8 morphological types have been reported (Gupta and Stopfer 2012).

**Fig. 6.**
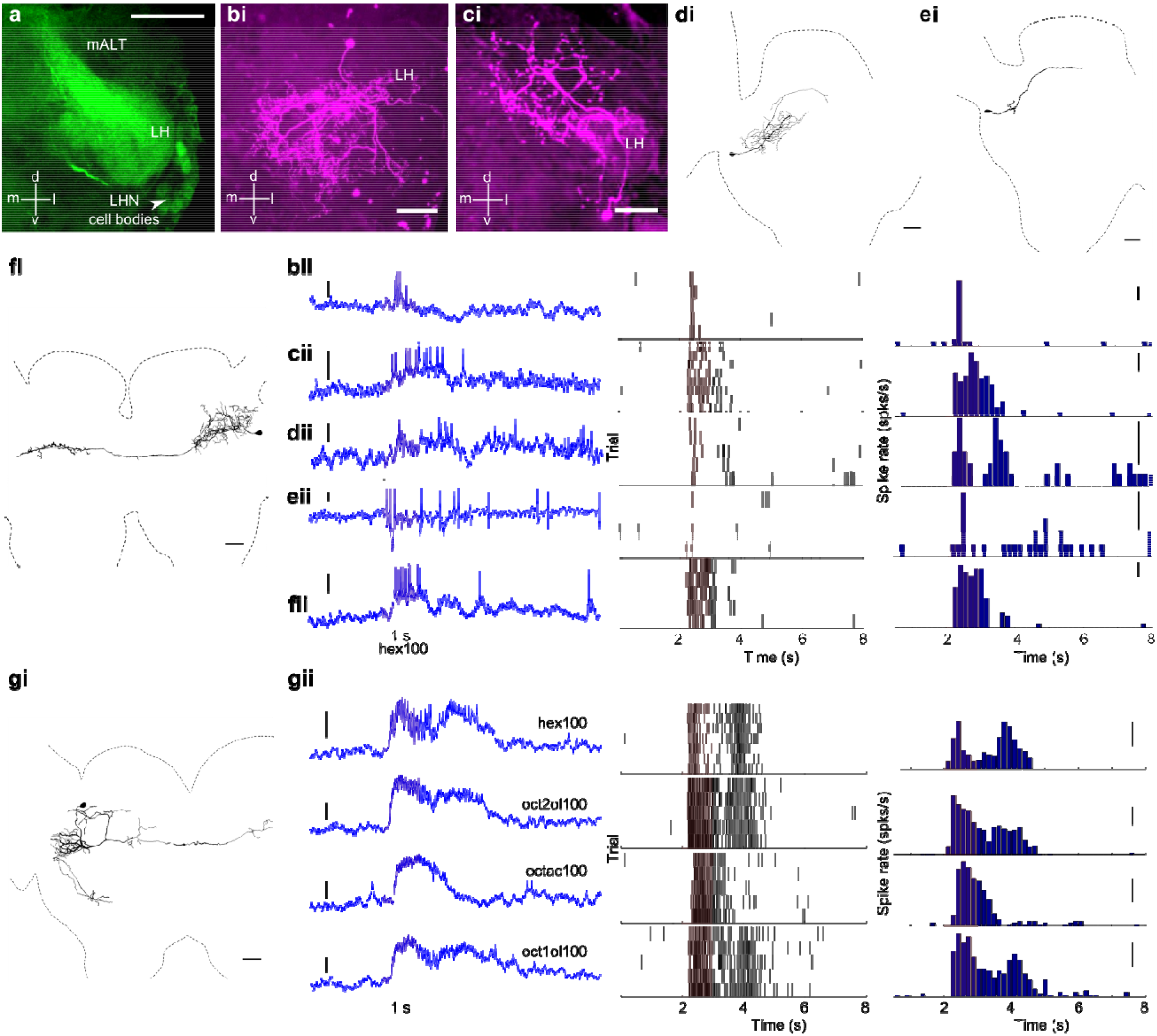
Morphology and odor response of lateral horn (LH) neurons in *H. banian*. **(a)** Dye fill from calyx of MB shows the cell bodies of LH neurons (arrowhead). Scale bar: 100 μm **(bi-gi)** Six distinct morphological types of LH neurons were filled in *H. banian*, with either sparse or very dense arborization in the LH area. **(bi)** and **(ci)** are z projections from a confocal stack while the rest **(di-gi)** are tracings made from the original confocal stack. Scale bar: 100 μm. The odor responses (raw traces, rasters and PSTH) of each of the filled LHNs are shown in **(bii-gii)**. Scale bar: Raw traces, **(bii)** 5 mV, **(cii-fii)** 2 mV, **(gii)** 5 mV; PSTH, **(bii-fii)** 5 spikes /sec, **(gii)** 20 spikes/sec

### Fourth order neurons-bLNs and GGN

Two distinct morphological types of MB extrinsic neurons arborizing in the β-lobe have been reported in *S. americana*-bLN1 and bLN2 (MacLeod et al. 1998; Gupta and Stopfer 2014). Both types were found in *H. banian*. bLN1 has its cell body near the midline of the brain and it has dense arborizations in the β-lobe and LH. It also extends sparse fibers in the superior protocerebral area (Fig. 7a). bLN1 was found to be GABA-negative in our immunohistochemical assay. We found two subtypes of bLN2 in *H. banian*-both types (Fig. 7c and 7e) extend their neurites to α-lobe, β-lobe and the peduncle of MB. In addition, both subtypes feed back to the MB. The difference between the two subtypes arises in the pattern of their innervation in the MB. One subtype has neurites which reach the MB calyx and spread there horizontally (Fig. 7c, arrowhead) while the other subtype has neurites which look restricted to the peduncular region in the calyx, most likely accessory calyx (Fig. 7e, arrowhead). All the three bLNs which we filled, showed spontaneous baseline activity and vigorous odor responses to all the odorants tested (Fig. 7b for bLN1, Fig. 7d for bLN2 subtype 1, and Fig.7f for bLN2 subtype 2). Mass dye fills from the β-lobe in *H. banian* (Fig. 7g) show a cluster of 13-16 bLN2 neurons, a number slightly higher than that reported in *S. americana* (Gupta and Stopfer 2014).

**Fig. 7.**
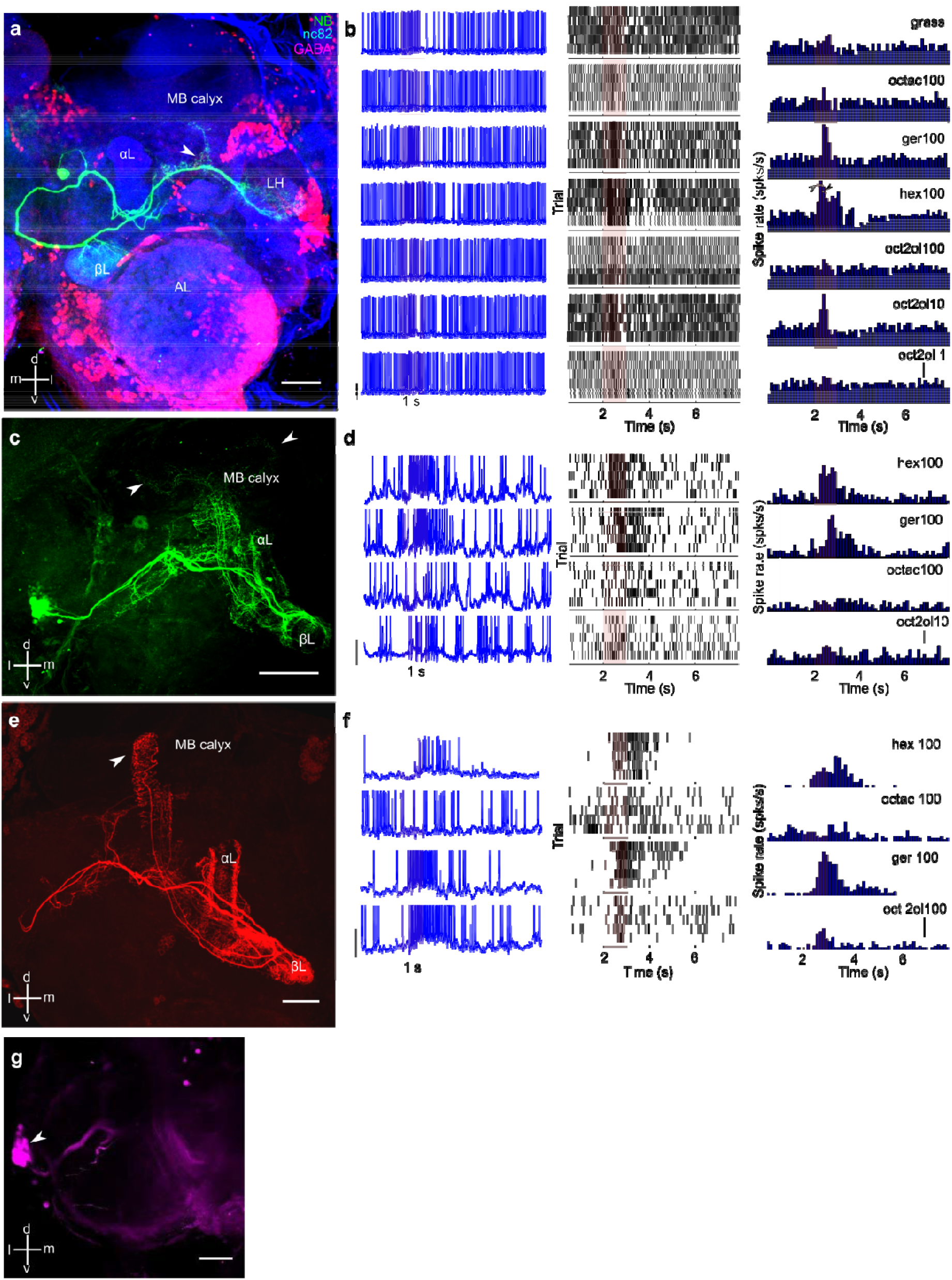
Morphology and odor response of β-lobe neurons (bLNs) in *H. banian*. **(a)** Triple-labelled intracellular fill of bLN1 with nc82 (in blue; to demarcate the synaptic densities) and GABA-positive elements (in magenta). The cell body is located near the midline of the brain; it has dense arborizations in the β-lobe of MB and LH and is not GABA-positive. A few projections are also seen in the superior protocerebral area (arrowhead). Scale bar: 100 μm **(b)** Odor response of bLN1 to different odorants shows that it responds to all of them, sometimes vigorously. Scale bar: 10 mV (raw traces); 20 spks/s (PSTH); open arrowhead ~130 spks/s, closed arrowhead ~60 spks/s **(c)** and **(e)** are intracellular fills of two subtypes of bLN2. Both of them innervate the peduncle, α-lobe and β-lobe of MB, with different patterns of arborization. One subtype **(c)** arborizes in the calyx of MB while the other **(e)** appears to innervate the accessory calyx. **(d)** and **(f)** represent the odor responses of the two subtypes to different odorants. They respond with different temporal patterns to the different odorants tested. Scale bar: 10 mV (raw traces); 20 spks/s (PSTH) **(g)** Mass dye fill from the β-lobe shows a cluster of 13-16 bLNs in the lateral protocerebral area (arrowhead). Scale bar: 100 μm

The cell body of GGN in *H. banian* is located ventral to the LH synaptic area and it has dense arborizations in the MB calyx, α-lobe and the LH (Fig. 8a). GGN is GABA-positive in *H. banian* (Fig. 8a, inset), similar to that in *S. americana* (Leitch and Laurent 1996; Papadopoulou et al. 2011). GABA immunohistochemistry on the whole brain shows that the GABA-positive arborization in the MB calyx of *H. banian* is separated into two distinct regions (Fig. 8b). We are not sure, if the source of the two layers of GABAergic arborization in the MB calyx is only from GGN or from two different sources, one of which is the GGN. In addition, dye fill from the calyx of MB in multiple preparations, has shown the existence of a single GGN-like fiber in the lateral protocerebrum, thus pointing to the fact that only one GGN is present in each half of *H. banian* brain (Fig. 8c). Intracellular recording of GGN shows the same response properties as in the other studied species, *S. americana* (Fig. 8d). The membrane potential consists of reliably detected IPSPs, which are thought to be coming from IG (Inhibitor of GGN), a neuron with unknown morphology (Papadopoulou et al. 2011). The odor response consists of depolarization of the membrane potential and this response is linearly scaled with log odor concentrations (Fig. 8d).

**Fig. 8.**
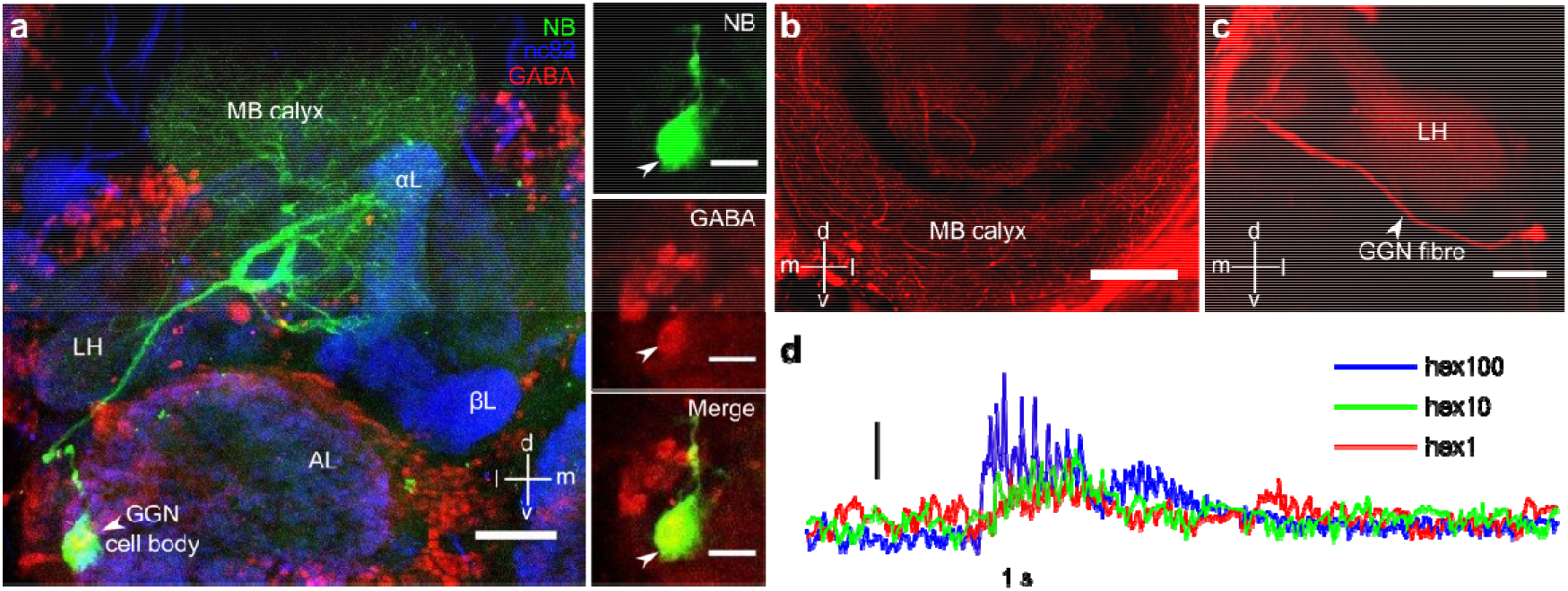
Morphology and odor response of giant GABAergic neuron (GGN) in *H. banian*. **(a)** Intracellular fill of GGN shows that it has dense and wide spread arborization in the calyx of MB. GGN also projects to α-lobe and LH. Anti-GABA immunohistochemistry on the same cell shows that it is GABAergic (inset). Scale bar: 100 μm; 50 μm for inset **(b)** Anti-GABA immunohistochemistry on the whole brain from another animal shows two distinct layer of GABAergic innervation in the calyx of MB. Scale bar: 100 μm **(c)** Mass dye fill from the calyx of MB shows a single-GGN like fiber originating from a single large somata in the lateral protocerebral lobe, implying that there is a single GGN in each half of the brain in *H. banian*. Scale bar: 50 μm **(d)** Odor response of GGN to different concentrations of an odorant. The depolarizing response of GGN increases in strength corresponding with increasing log concentration of odorant. Scale bar: 10 mV

### A novel type of MB extrinsic neuron (MBEN)

We recorded and filled three different subtypes of MBENs, whose cell bodies are located in the superior lateral protocerebral lobe. These have not been reported till now, to the best of our knowledge, in the olfactory pathway. All the three neurons innervate bilaterally (fills from three different animals). One subtype which we filled (Fig. 9a) has dense arborizations in both the alpha lobes of MB. This neuron tested negative for anti-GABA antibody and showed very weak response to different odorant stimuli (Fig. 9c). The second subtype (Fig. 9b) has dense arborizations in the ipsilateral peduncle of MB and sparse arborization in the contralateral peduncle. This neuron was GABA-positive and showed excitatory response to odorants while maintaining a consistent baseline rate (Fig. 9d). The third MBEN which we filled, does not seem to arborize in the ipsilateral half, but crosses the midline and arborizes sparsely in the protocerebral area (Fig. 9e). This neuron was weakly responsive to the odorant 1-hexanol (Fig. 9f). Even though all three of them arborize in the output regions of the third order neurons in the olfactory pathway, they show weak response to odor stimuli. This is in contrast to the vigorous odor responses seen in all the known fourth order neurons like bLNs and GGN in the olfactory pathway.

**Fig. 9.**
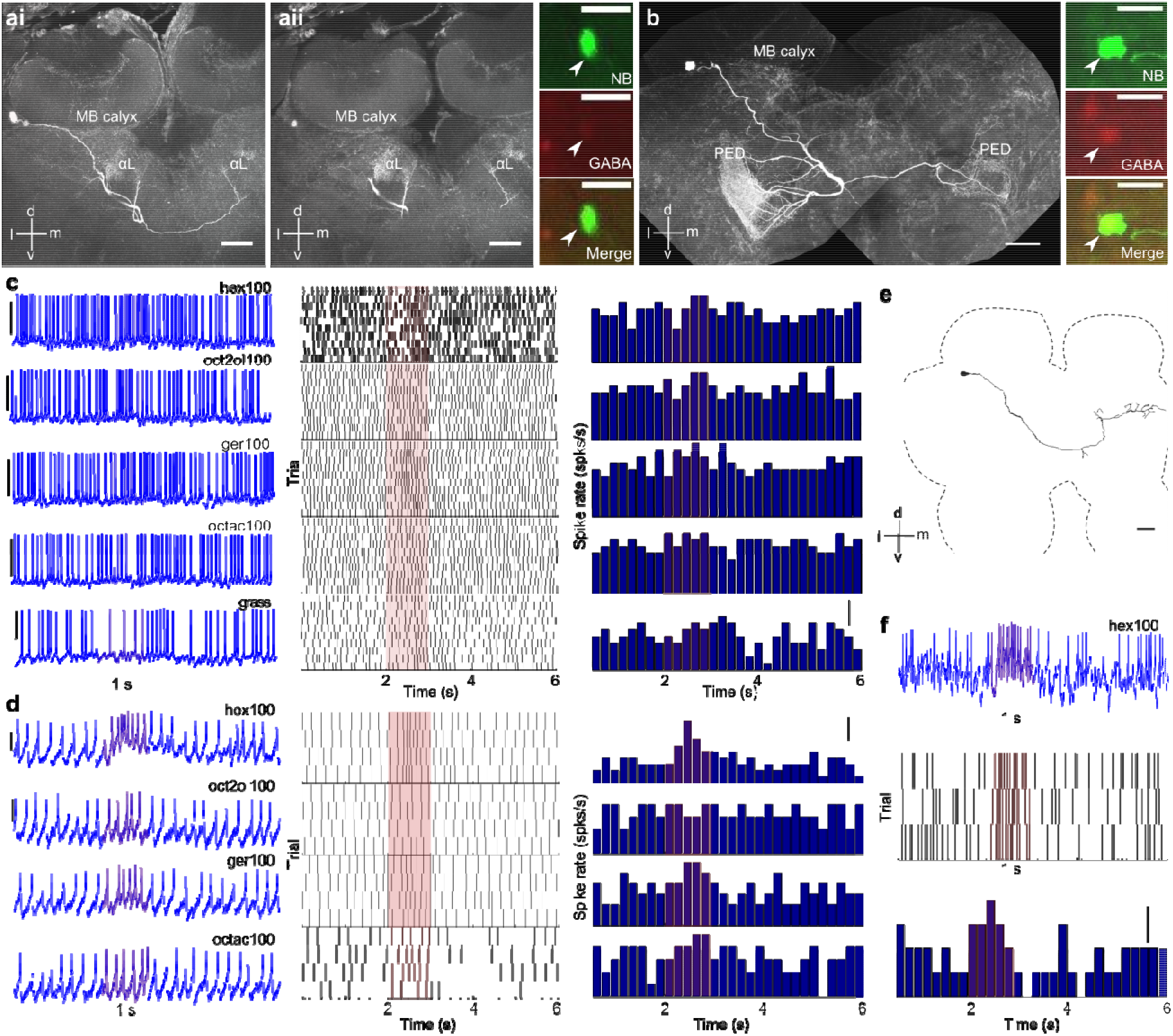
Morphology and odor response of a newly discovered set of bilateral MB extrinsic neuron (MBEN). **(ai** and **ii)** A bilateral MBEN (type 1) whose cell body is located in the superior lateral protocerebral (SLPL) area and it has dense arborizations in both the α-lobes. Anti-GABA immunohistochemistry shows it to be non-GABAergic (inset, arrowhead) Scale bar: 100 μm; inset 50 μm **(b)** Another bilateral MBEN (type 2) with its cell body at the SLPL, and dense innervation in the peduncle of MB on the ipsilateral side. The neuron crosses the midline below the central complex and projects to the contralateral peduncle of MB with sparse innervation. The neuron is GABAergic. (inset, arrowhead) Scale bar: 100 μm; inset: 50 μm **(c)** Odor response of the type 1 MBEN to five different odorants. The neuron had very little response to odorants though it innervates downstream of Kenyon cells densely. Scale bar: 0.01 mV (raw traces); 5mV (grass, raw trace); 2 spks/s (PSTH) **(d)** Weak odor response of type 2 MBEN to four odorants. Scale bar: 2 mV (raw traces); 2 spks/s (PSTH) **(e)** and **(f)** show the morphology and odor response of a third type of MBEN. This neuron also has its cell body in the SLPL area and it crosses the midline to project to the contralateral protocerebral lobe. Its odor response consists of slight increase in firing rate. Scale bar: 100 μm; 2 μV (raw trace), 2 spks/s (PSTH)

**Fig. 10.**
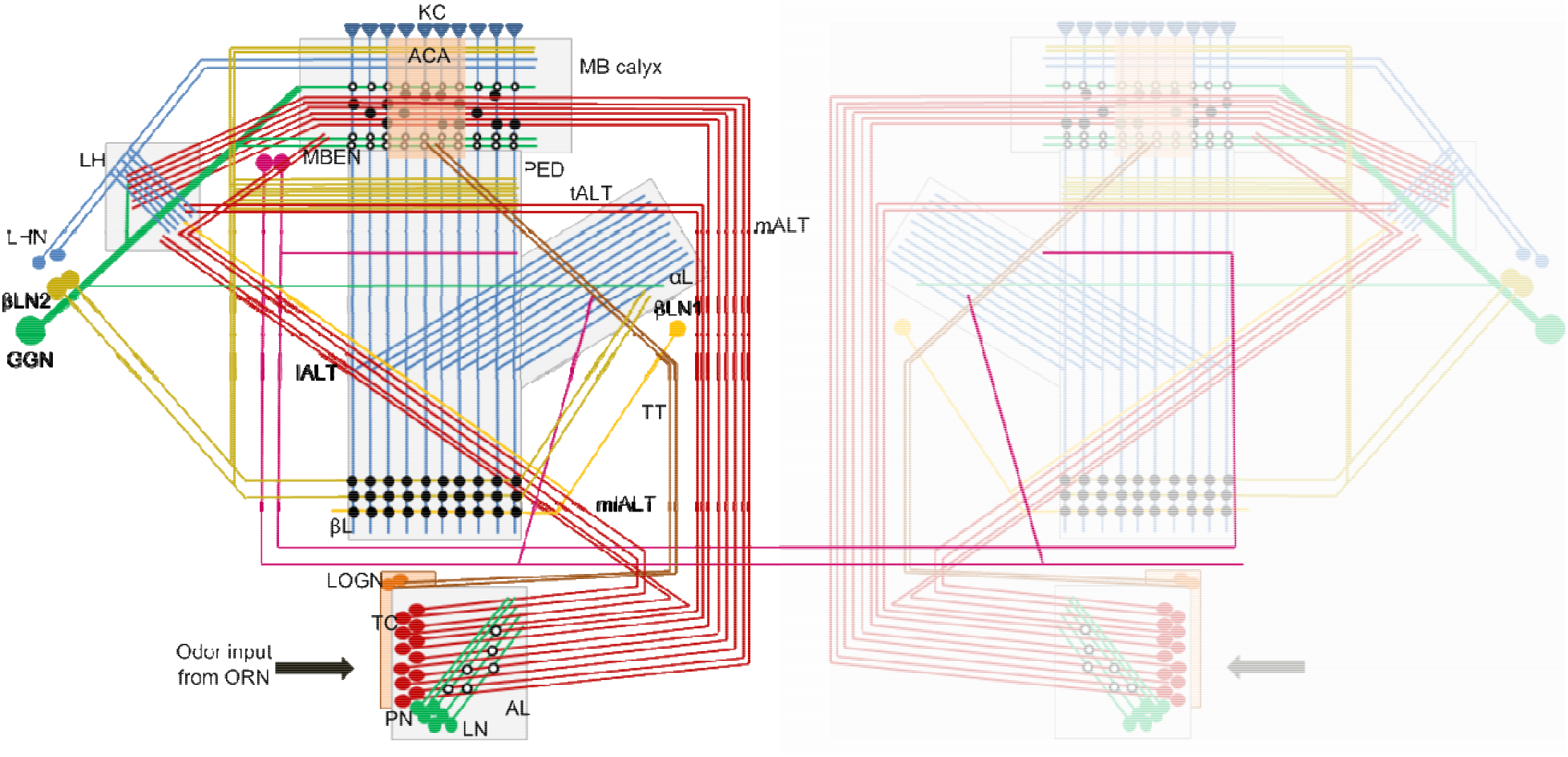
Schematic overview (modelled on published data in *Schistocerca spp.* and primary data in this study) of the olfactory circuit in *H. banian* from the second order (antennal lobe) to the fourth order (β-lobe and GGN). The olfactory circuit in *H. banian* begins with the first order ORNs, housed in sensilla on the antenna. The ORNs make synapses on the PNs and LNs present in the second relay center, the AL. The PNs are the only output from AL and they relay the olfactory information through multiple tracts (mALT, lALT, mlALT and tALT) to the KCs in MB and LH, the third order centers. The axons of the KCs form the peduncle of the MB which bifurcates into the α- and β-lobes. The KCs make synapse with the fourth order neurons in the β-lobe (bLNs) and with the GGN and bLNs in the α-lobe. The Giant GABAergic neuron (GGN) has neurites in the MB calyx, α-lobe and LH. The tritocerebral tract, part of the gustatory circuit, arises from the cell bodies of LOG located dorso-laterally near the AL and terminates in the accessory calyx of MB. Newly discovered bilateral MBENs in *H. banian* have their cell bodies near the superior lateral protocerebral lobe and they project either to α-lobes or peduncles in both halves of the brain. ORN: olfactory receptor neuron; AL: antennal lobe; LN: local neuron; PN: projection neuron; mALT: medial antennal lobe tract; mlALT: mediolateral ALT; tALT: transverse ALT; lALT: lateral ALT; GGN: giant GABAergic neuron; LH: lateral horn; MBEN: mushroom body extrinsic neuron; MB calyx: mushroom body calyx; PED: peduncle; KC: Kenyon cell; bLN: β-lobe neuron; αL: α-lobe; βL: β-lobe; TT: tritocerebral tract; ACA: accessory calyx; LOG: lobus glomerulus; LOGN: lobus glomerulus neurons; TC: tritocerebrum; open circles: inhibitory synapse; closed circles: excitatory synapse

## Discussion

A major goal of the present study was to investigate the differences and similarities of neurons involved in olfactory processing at different levels (from second order to fourth order; morphologically as well as physiologically) between two Orthopteran species, *Hieroglyphus banian* and *Schistocerca americana* belonging to different sub-families (Hemiacridinae and Cyrtacanthacridinae respectively). Our results from *H. banian* when compared with the existing literature in *S. americana* show that the olfactory circuit is highly conserved till the fourth order neurons even though the two sub-families diverged ~57 million years ago (Song et al. 2018). We believe this to be the first study that compares from the second order up to the fourth order neurons in a sensory system across subfamilies among any insect.

Apart from comparing the olfactory circuits of *H. banian* and *S. americana*, and the fact that all the results in this study have been shown for the first time for the species *H. banian,* we present the following new findings with respect to the order Orthoptera which have not been reported before: (1) three new tracts from antennal lobe to mushroom body (MB) and a new tract from the lateral horn to the MB, (2) the LFP measured from the MB cell body layer and primary calyx during odor response are negatively correlated, (3) during odor response, the power of lower frequencies present in the LFP recorded from the MB calyx decreases while the higher frequency component of the LFP increases in power over the course of trials, (4) five new morphological types of lateral horn neurons (LHNs), different from those reported in *Schistocerca americana* (Gupta and Stopfer 2012), (5) a new subtype of β-lobe neuron type 2 (bLN2), (6) a novel bilateral MB extrinsic neuron (MBEN).

### Rationale behind the methodology used to establish similarities between the two species

To the best of our knowledge, this is a first study which compares the olfactory circuit from the second order to the fourth order and shows that the olfactory circuit elements at the higher levels are highly conserved in terms of morphology and physiology. The use of intracellular recordings, intracellular dye fills and immunohistochemistry to compare neurons is a well-established one (Römer et al. 1988, where the morphology and soma position of the auditory neurons in locusts and bush crickets were used to establish homology between them). In *Drosophila*, neural labelling under genetic control is used extensively for anatomical study (Wong et al. 2002; Lai et al. 2008; Tanaka et al. 2012a). On the other hand, some purists consider it imperative to do a lineage analysis of the neurons to establish that they are homologous in nature (Boyan 1993). But it might not be practical to do lineage analysis or neural labelling under genetic control in all cases-and numerous studies across insect species have shown the efficacy and adequateness of comparing neurons using intracellular recordings and dye fills (Rössler and Zube 2011; Namiki and Kanzaki 2011; Tanaka et al. 2012b). In *Drosophila*, new tracts from AL to MB were conveniently discovered using mass fills from AL, in addition to those discovered by labelling neurons under genetic control (Tanaka et al. 2012b). Mass fills were also used to discover new AL tracts in the heliothine moth, *Heliothis virescens* and compare it to other moths (Ian et al. 2016). Thus, the use of these techniques to compare the olfactory elements in any two species is well-established.

### Variation and adaptation of the olfactory circuit at the peripheral level versus higher level

Variability in the morphological architecture at the peripheral level is observed across insect species because of the selection pressures acting on it while encountering diverse problems-ecological or functional (Hansson and Stensmyr 2011). Within Orthoptera, there is a wide diversity at the peripheral level (AL), in terms of numbers of glomeruli (as few as 12 to as large as 1000s of glomeruli), types of glomeruli (unique/microglomeruli), arborization patterns of receptor neurons and projection neurons (from uni-, bi-to multiglomerular PNs; Ignell et al. 2001). In Lepidoptera (moths), Coleoptera (beetles), Hymenoptera (ants) and Diptera (*Drosophila*) too, the glomerular counts in the antennal lobe varies across species (Kondoh et al. 2003; Kelber et al. 2009; Bisch-Knaden et al. 2012; Kollmann et al. 2016). One limitation of this study is that we have not numerically characterized the glomeruli numbers in AL or the numbers of ORNs, AL PNs, AL LNs or MB KCs in *H. banian*, but we have found that they are conserved broadly in terms of the nature of their physiology, morphology and immunohistochemical character when compared to *S. americana*. The neurons of the third and fourth levels (bLNs, GGN and LHNs) are quantitatively, morphologically and physiologically similar in both the species.

### Variation in lifestyle and habitat may not necessarily translate into variation in the olfactory circuit

*H. banian* is a species endemic to the Indian subcontinent and Vietnam whereas *S. americana* is endemic to North America (Cigliano et al. 2018). They differ in their food preferences too (Capinera 1993; Squitier and Capinera 1996; Das et al. 2002; Mandal et al. 2007). Considering the varying habitats which they occupy and the differences in host plant preferences, we expected that since they would encounter and solve dissimilar ecological problems during their life histories, it would give rise to some variability in their olfactory circuit in terms of morphology or physiology. But our results from *H. banian,* compared with the literature on *S. americana,* show that the two species seem to be highly conserved at the higher levels, even though they belong to different sub-families. This may be due to the stabilizing selective forces at work to prevent major adaptations in the circuit at the higher levels. On the other hand, minor differences at the peripheral level between species are expected as the selective forces might act on the initial olfactory circuit to develop specific adaptations enabling the system to solve problems specific to food habits and habitats.

### Olfactory circuit of *H. banian* is similar in organization to that of *S. americana* with few differences

The antennae of *H. banian* show all the four types of sensilla (basiconica, trichodea, coeloconica and chaetica) previously reported in *S. gregaria* (Ochieng et al. 1998).

There is only one reported AL tract in *S. americana* or its related species *S. gregaria*-the mALT (Leitch and Laurent 1996; Hansson and Anton 2000; Ignell et al. 2001; Anton et al. 2002; Galizia and Rössler 2010). Another orthopteran, *Locusta migratoria* is also reported to have a single AL tract to MB-the mALT (Ernst et al. 1977). *Tetrix subulata* (Family: Tetrigidae) is the only orthopteran species which has been reported to have two AL tracts-the mALT and the lALT (Ignell et al. 2001). In *H. banian*, on the other hand, dye fills from the AL in the present study, have shown additional tracts, the medio-lateral ALT (mlALT), the lateral ALT (lALT) and the transverse ALT (tALT), which correspond to those reported in Lepidoptera, Hymenoptera, Diptera and Blattodea (Galizia and Rössler 2010; Tanaka et al. 2012b, a; Ian et al. 2016). The probable reason for not discovering these additional tracts might be their selective staining as they seem to have few fibers running through them. In the present study too, we did not see all the tracts in all the successful fills which we had. If we go by Ignell et al. (2001), then the lateral most tract in our fill is the lALT. But the origin of this tract is medial to that of the mlALT in *H. banian*, and it projects only to the LH. This is in contrast to the trajectory of the lALT found in *Apis mellifera* (Kirschner et al. 2006), other hymenopterans (Rössler and Zube 2011) and the moth, *Heliothis virescens* (Ian et al. 2016), where it first projects to the LH and then runs further and terminates in the MB. In *H. banian*, this trajectory is followed by the mlALT. There is one major distinction between the lALT of Hymenoptera and Lepidoptera and the mlALT in *H. banian*. In Hymenoptera, the lALT forms one half of a dual parallel pathway between AL and MB. On the other hand, the lALT in Lepidoptera and the mlALT in *H. banian* has fewer fibers as compared to the mALT, which forms the major tract to the MB (Kirschner et al. 2006; Galizia and Rössler 2010; Martin et al. 2011; Rössler and Zube 2011).

At the subfamily level, if we compare *Apis mellifera* (subfamily: Apinae) and *Bombus terrestris* (subfamily: Bombinae), 5 tracts are known in *A. mellifera* (Kirschner et al. 2006) while only two AL tracts have been reported in *Bombus spp.*-mALT and lALT (Strube-Bloss et al. 2015). The carpenter ant, *Camponotus floridanus* (subfamily: Formicinae), *Harpegnathos saltator* (subfamily: Ponerinae) and *Atta vollenweideri* (subfamily: Myrmicinae) all have 5 tracts (Zube et al. 2008; Rössler and Zube 2011).

In Lepidoptera (moths), *Manduca sexta* (subfamily: Sphinginae; Homberg et al. 1988) *Heliothis virescens* (subfamily: Heliothinae; Ian et al. 2016) have 5 AL tracts while only 3 have been shown for *Bombyx mori* (subfamily: Bombycinae; Kanzaki et al. 2003; Seki et al. 2005). In *Periplaneta americana* too (order: Blattodea), 5 AL tracts are found (Malun et al. 1993) while in the Dipteran, *Drosophila melanogaster,* 4 have been found (Stocker et al. 1990; Tanaka et al. 2012b; Ito et al. 2014).

What does it mean to have multiple tracts? Does it point to a more complex circuit? Galizia and Rossler (2010) reported the presence of a single AL tract (mALT) in primitive insects like basal Apterygota, the Archaeognatha (bristletails; *Malachis germanica*) which have no MBs and in Zygentoma (silverfish). Basal holometabolus insects of the order Coleoptera (beetles) also have a single AL tract from AL to MB (Galizia and Rössler 2010). According to them, Acrididae insects have a single tract from AL to MB (mALT) because all the known PNs are multiglomerular in this family. The presence of multiple tracts points to higher complexity in the kinds of PNs present (uniglomerular or multiglomerular) or to different channels for olfactory information processing (Galizia and Rossler 2010). In *Manduca sexta* and *Heliothis virescens,* the mlALT is composed of only mPNs while the lALT and mALT have both u- and mPNs (Homberg et al. 1988; Helge et al 2007; Ian et al. 2016). Many PNs of the mlALT in both *M. sexta* and *H. virescens* have also been shown to be GABAergic (Hoskins et al. 1986; Berg et al. 2009). Similar is the case in Hymenoptera (bees and ants), where uPNs project through lALT and mALT to higher olfactory centers while mPNs comprise the three mlALTs (Kirschner et al. 2006; Zube et al. 2008; Rössler and Zube 2011). In bees, it is known that uPNs belonging to different tracts differ in their physiological properties and may process different features of odor information (Abel et al. 2001; Müller et al. 2002; Peele et al. 2006; Kirschner et al. 2006; Galizia and Rössler 2010). Even the neurotransmitter is different for PNs of lALT and mALT in honey bees. The PNs of only mALT are cholinergic (Kreissl and Bicker 1989) while those of lALT show strong taurine-immunoreactivity (Schäfer et al. 1988; Kreissl and Bicker 1989). Few GABAergic PNs are also found in mALT and mlALT (Schäfer and Bicker 1986). The discovery of multiple AL tracts in *H. banian* may thus probably imply that that there might be a possibility of finding uPNs in this species, or that there could be different clusters of PNs like those found in Lepidoptera and Hymenoptera, or that there could be different channels for processing different kinds of information like that found in Hymenoptera.

The tritocerebral tract (TT) in *H. banian,* which arises from axons of LOG cell bodies and terminates in the accessory calyx is similar to the one shown in *Schistocerca gregaria* (Homberg et al. 2004) and *Locusta migratoria* (Ernst et al. 1977). The TT has not been shown in *S. americana* as of now. Though the TT shown in *S. gregaria* was visualized by using antibody against Mas-allatotropin and our tract was visualized by direct dye injection in the LOG cell body area, the trajectories followed by both of them to the MB accessory calyx are very similar. According to an exhaustive study across insect orders by Farris (2008), TT or SCT (sub-oesophageal calycal tract, as it is referred to in some insects) is ubiquitous in insects. The termination site of the TT may, however, vary across insect species depending on the presence or absence of the accessory calyx in MB (Farris 2008).

We have also discovered a new tract, the CT (curved tract) running between the LH and the MB calyx. This tract has not been reported in any insect species till now, as far as we know and further studies would be required to find its role in the circuit.

*H. banian* also has microglomerular organisation in the AL like *S. americana* (Laurent and Naraghi 1994). The number of microglomeruli in the AL was not quantified for this species. In fact, microglomerular organization of the AL is found across species in the family Acrididae (*Locusta migratoria*,Ernst et al. 1977; *S. gregaria*,Anton and Hansson 1996; Ignell et al. 2001) and is considered to represent a higher complexity in organization. In line with this, the AL in lower Orthoptera and its sister order Blattodea (*Periplaneta americana*) which have unique glomeruli are considered to be primitive in nature (Ignell et al. 2001). An exhaustive study across 22 different families in Coleoptera (beetles) also shows a wide diversity in AL glomerular organization which is more or less conserved at the family level (Kollmann et al. 2016).

The PNs and LNs which we filled and recorded in this study look similar physiologically and morphologically to those found in *S. americana* (Laurent and Naraghi 1994; MacLeod and Laurent 1996) and *S. gregaria* (Anton and Hansson 1996; Ignell et al. 2001). The PN which we filled in *Hieroglyphus* is multiglomerular, projects to a large area in the MB calyx and shows spatio-temporal coding in its odor response like the PNs in *S. americana* (Laurent and Davidowitz 1994; Laurent and Naraghi 1994; Laurent 1996; Wehr and Laurent 1996; Anton et al. 2002; Perez-Orive et al. 2002; Mazor and Laurent 2005). But with the discovery of new AL tracts in *H. banian*, there arises a need for a comprehensive investigation of PN types. The PNs show Na^+^ spikes (Distler 1990) whereas the LNs show short spikes putatively calcium-mediated (Laurent and Davidowitz 1994; Laurent 1996). This is consistent with that found in *S. americana.* In *Schistocerca*, the PN has been shown to be excitatory and GABA-negative (Leitch and Laurent 1996). *Manduca sexta* and *Bombyx mori* which belong to different subfamilies (Sphinginae and Bombycinae respectively) also have similar morphological types of PNs (Namiki and Kanzaki 2011). The LN in *H. banian* is morphologically similar and GABA-positive like the one found in *S. americana* (MacLeod and Laurent 1996).

### Higher order olfactory centers are conserved between *H. banian* and *S. americana*

The LFP oscillations observed during odor response from the MB calyx in *H. banian* show a predominant frequency of ~25 Hz which is similar to that reported in *S. americana* (Laurent and Naraghi 1994; Laurent 1996). We could not find a comparison of LFP oscillation frequency at subfamily level in any other insect order. However, known LFP frequencies vary widely across insect orders from ~10 Hz in *Drosophila* (Tanaka et al. 2009), ~17 Hz in the wasp, *Polistes fuscatus* (Stopfer et al. 1999), to ~35 Hz in the moth *Manduca sexta* (Ito et al. 2009). We do not know if the variability in LFP frequency has some significance for information processing. We also do not know the exact cause for this variation though it is likely to do with the strength, timescale and number of synapses in the antennal lobe. We have showed for the first time that the oscillations in LFP recorded from the cell body layer of the MB during odor response is out of phase and negatively correlated with that recorded from the MB calyx. This is consistent with the anatomical structure of the MB where the KC cell bodies and the synaptic regions are anatomically segregated in adjacent areas. This has implications for interpreting phase relationships recorded intracellularly from neurons in different areas and MB LFP (calyx or cell body layer). We also show that during odor response, the lower frequency component of LFP decreases while the higher frequency component increases in power. The reason for this change and its significance in olfactory coding is not clear yet. In olfactory system of mammals, odor evoked oscillations recorded in vivo range from 15-35 Hz to 40-90 Hz (Buonviso et al. 2003; Martin et al. 2004).

We could not fill any Kenyon cell in our preparations, but the recordings which we got from them, show characteristics of KC physiology-low baseline rate and sub-threshold membrane oscillations during odor response. This corresponds well to the KC physiology in *S. americana* (Laurent and Naraghi 1994; Perez-Orive et al. 2002; Broome et al. 2006; Jortner et al. 2007).

The LHNs, which we recorded and filled from *H. banian,* show a wide variety in morphology and response pattern to odor stimuli. Of all the LHN fills in *H. banian*, only one fill corresponded to one of the types (named C3) reported by Gupta and Stopfer (2012)in *S. americana*. In principle, we can say that there is wide diversity in the morphological types of LHNs in both the species and more rigorous study might be required to elucidate the various LHN types and the function which they might play in olfactory information processing. The type of LHN which feeds back to the MB, one of the types shown by Gupta and Stopfer (2012)has also been shown in *Bombyx mori* (Namiki et al. 2013). There is no study, as of now, which compares the LHNs’ morphology and physiology between species diverging at the subfamily level. Apart from grasshopper and locusts, LHNs have been studied only in *D. melanogaster* (Fişek and Wilson 2014). *D. melanogaster* has two groups of LHNs-type I and type II, distinguished by spatially separated cell clusters. The LH has been implicated to play a role in mediating innate behavior, multimodal integration, bilateral coding and concentration coding in *S. americana* (Gupta and Stopfer 2012). A wide range of roles have been attributed to the LHNs across insect species (review by Schultzhaus et al. 2017). Recently, Chin et al. 2018 reported that LH plays a role in negative oviposition behavior in *D. melanogaster*. We did not investigate the multimodality of the LH neurons in this study.

Similar to those found in *S. americana* (MacLeod et al. 1998; Cassenaer and Laurent 2007; Gupta and Stopfer 2014) two types of β-lobe neurons (bLNs) were found in *H. banian*, type 1 (1 neuron) and a cluster of type 2 bLNs. We counted the number of bLN2s and found that it is slightly more than that reported for *S. americana*. We also propose that there may be two subtypes of bLN2, one which feeds back to the MB calyx and the other which arborizes in the peduncle and the accessory calyx of MB. Apart from *S. americana*, output neurons of MB have been studied only in the species *Apis mellifera* (α-lobe neurons; Bicker et al. 1985; Rybak and Menzel 1993, 1998), *Periplaneta americana* (Li and Strausfeld 1997; Okada et al. 1999), *Acheta domesticus* (Schildberger 1984) and *Drosophila melanogaster* (Tanaka et al. 2008; Aso et al. 2014).

The GGN, in *H. banian*, shows striking similarities in morphology and physiology to that in *S. americana* (Leitch and Laurent 1996; Papadopoulou et al. 2011) and *S. gregaria* (Homberg et al. 2004). All three species have arborizations of GGN in the MB calyx, α-lobe and the LH. Odor response of GGN in both *H. banian* and *S. americana* (Papadopoulou et al. 2011; Gupta and Stopfer 2012, 2014) show graded potentials with clear IPSPs. Both of them are immunopositive for the inhibitory neurotransmitter GABA (Leitch and Laurent 1996). Except APL (anterior paired neuron) in *Drosophila* (Yasuyama et al. 2002; Tanaka et al. 2008; Liu and Davis 2009; Papadopoulou et al. 2011), other insects *Periplaneta americana* (Weiss 1974; Yamazaki et al. 1998; Takahashi et al. 2017), *A. mellifera* (Bicker et al. 1985; Rybak and Menzel 1993; Grünewald 1999) and *Manduca sexta* (Homberg et al. 1987) have multiple GGN-like neurons (4, ~50, ~150 respectively). The GGN-like neurons are called Calycal Giants (CGs) in *Periplaneta* and include 3 spiking and one non-spiking neuron. All four of them are GABAergic and feed back to the MB calyx. Only the non-spiking CG looks similar to the GGN in morphology and physiology (Takahashi et al. 2017). The zonation in GABAergic innervation in the MB, which we see in Fig. 8b, has also been reported by Gupta and Stopfer (2012). They observed GABAergic fibers of unknown origin innervating the accessory calyx of MB in *S. americana*. Takahashi et al (2017)have also reported zonation by the 4 CGs in their innervation of the MB calyx. The zonation by CGs in the MB calyx of *Periplaneta* was attributed to the innervation pattern by the two groups of uPNs in MB calyx (Takahashi et al. 2017).

Here, in this study, we are reporting for the first time a cluster of bilateral neurons whose cell bodies are located near the edge of the MB calyx, in the superior lateral protocerebral area and project to higher olfactory center-the MB. These neurons, three of which we filled and recorded from, show widely varying morphologies but weak odor response. Such neurons have not been reported in *S. americana* yet and more studies are required in future to dissect their role in the olfactory circuit.

### How does the olfactory circuit compare among vertebrates diverging at the subfamily level?

From the available literature, the olfactory circuit of mouse (subfamily: Murinae) and humans (subfamily: Homininae) can be compared to see their similarities and dissimilarities as an example of vertebrates diverging at the subfamily level (Sinakevitch et al. 2018). At the peripheral level, a few differences are there-for example, there are fewer olfactory receptors in humans (~350; Glusman et al. 2001; Maresh et al. 2008) than mouse (~1200; Zhang et al. 2007) and each olfactory receptor neuron arborizes in ~16 glomeruli in humans (Maresh et al. 2008) as compared to ~2-3 glomeruli in mouse (Sinakevitch et al. 2018). Mouse olfactory bulb (OB) is also characterized by the presence of the sex specific vomeronasal organ as part of it. Vomeronasal organ is absent in human OB. There are differences in total numbers of various neuronal types (mitral, tufted and local neurons) in the OB between human and mouse but in principle the basic plan seems to be similar, not only between them but across mammals (Sinakevitch et al. 2018). Differences at higher levels are less understood in vertebrates because there are very few studies dealing with higher order neurons.

## Conclusion

In conclusion, the olfactory circuit in *H. banian* is very similar to that in *S. americana* from the second order to the fourth order, both in physiology and anatomy, even though they belong to different subfamilies, which diverged approximately 57 million years ago. Though future studies with emphasis on quantitatively characterizing the peripheral levels, AL and MB neurons and studying their immunohistochemical properties, in depth, would be required to completely substantiate any claims about similarity of the two species, the present study does throw some light on this with respect to the higher levels. This study can work in tandem and as a nice complement to develop *H. banian* as a tractable model organism like *S. americana,* for studying questions regarding information processing in a simple olfactory circuit.

## Supplementary figure

**Fig. A1.**
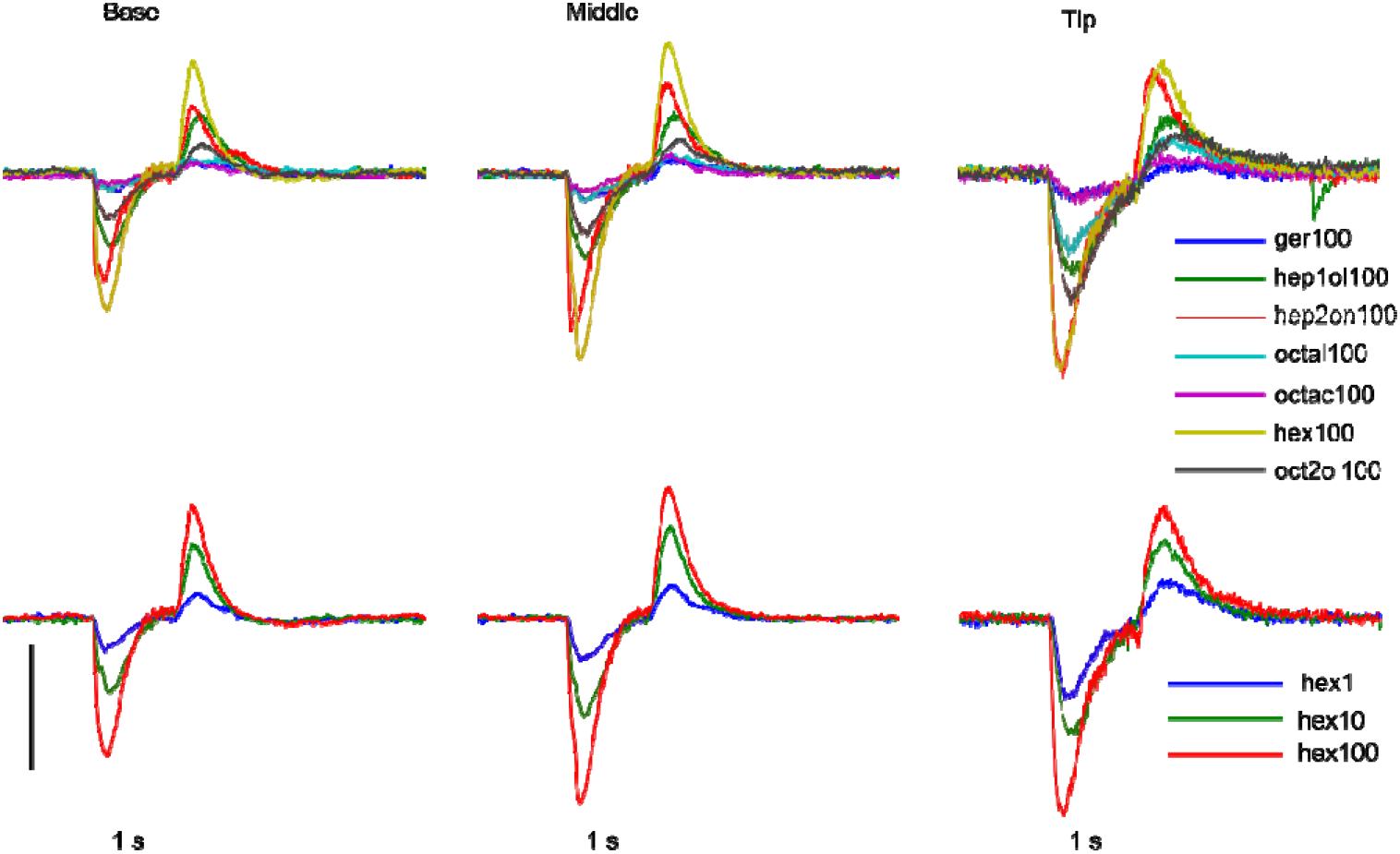
Electroantennogram (EAG) recorded from the antenna varies across its length in *H. banian.* EAG measured from the basal, middle and tip segments of the antenna of the same animal shows that there is difference in the population odor response of the sensilla present in those segments. The strongest response was recorded from the tip and the weakest from the basal segment of the antenna. This correlates nicely with the density of sensilla on the antennae, which is highest at the tip. 1-Hexanol (hex), 2-Octanol (oct2ol), Octanoic acid (octac), Geraniol (ger), Octanal (octal), 1-Heptanol (hep1ol), 2-Heptanone (hep2on) Scale bar: 300 μV

### Electroantennogram (EAG) recording method

EAG from the tip (distal two segments), middle and basal (proximal two segments) portions of the antennae was recorded using custom made blunt borosilicate glass microelectrodes (impedance <10MΩ after filling with saline). Reference electrode was Ag-AgCl wire inserted in the eye. The signal was amplified using Axopatch 200B (Axon Instruments) amplifier and band-pass filtered at 0.7 Hz-300 Hz. Subsequently, it was acquired at a sampling rate of 15 kHz using National Instruments USB acquisition system. The data was analyzed offline using custom made programme in MATLAB (Mathworks).

## Acknowledgements

We would like to thank the Center for Nanotechnology (Confocal Laser Scanning Microscope facility), Central instrumentation Laboratory (CIL, Confocal Laser Scanning Microscope facility) and School of Physics (FE-SEM facility), all at the University of Hyderabad for providing access to their infrastructure. We are grateful to Ms Nalini Manthapuram, Mr. M Prasad and Ms Sunitha for their expert technical assistance in using the CLSM and FE-SEM facilities respectively.

## Abbreviations

*ACA*: Accessory calyx
*AL*: Antennal lobe
*bLN*: β-lobe neuron
*CT*: Curved tract
*FE-SEM*: Field emission scanning electron microscopy
*GGN*: Giant GABAergic neuron
*IG*: Inhibitor of GGN
*KC*: Kenyon cell
*lALT*: Lateral antennal lobe tract
*LH*: Lateral horn
*LHN*: Lateral horn neuron
*LOG*: Lobus glomerulus
*LFP*: Local field potential
*LN*: Local neuron
*mALT*: Medial antennal lobe tract
*mlALT*: Medio-lateral antennal lobe tract
mPN: Multiglomerular projection neuron
*MB*: Mushroom body
*MBEN*: Mushroom body extrinsic neuron
*OR*: Olfactory receptor
*ORN*: Olfactory receptor neuron
*PSTH*: Peri-stimulus time histogram
*PN*: Projection neuron
*tALT*: Transverse antennal lobe tract
*TT*: Tritocerebral tract
uPN: Uniglomerular projection neuron

